# A vector-based strategy for olfactory navigation in *Drosophila*

**DOI:** 10.1101/2025.02.15.638426

**Authors:** Andrew F. Siliciano, Sun Minni, Chad Morton, Charles K. Dowell, Noelle B. Eghbali, Juliana Y. Rhee, L.F. Abbott, Vanessa Ruta

## Abstract

Odors serve as essential cues for navigation. Although tracking an odor plume has been modeled as a reflexive process, it remains unclear whether animals can use memories of their past odor encounters to infer the spatial structure of their chemical environment or their location within it. Here we developed a virtual-reality olfactory paradigm that allows head-fixed *Drosophila* to navigate structured chemical landscapes, offering insight into how memory mechanisms shape their navigational strategies. We found that flies track an appetitive odor corridor by following its boundary, alternating between rapid counterturns to exit the plume and directed returns to its edge. Using a combination of behavioral modeling, functional calcium imaging, and neural perturbations, we demonstrate that this ‘edge-tracking’ strategy relies on vector-based computations within the *Drosophila* central complex in which flies store and dynamically update memories of the direction to return them to the plume’s boundary. Consistent with this, we find that FC2 neurons within the fan-shaped body, which encode a fly’s navigational goal, signal the direction back to the odor boundary when flies are outside the plume. Together, our studies suggest that flies leverage the plume’s boundary as a dynamic landmark to guide their navigation, analogous to the memory-based strategies other insects use for long-distance migration or homing to their nests. Plume tracking thus uses components of a conserved navigational toolkit, enabling flies to use memory mechanisms to navigate through a complex shifting chemical landscape.

---

Navigating within a complex environment is a ubiquitous requirement for all animals, driving the evolution of diverse behavioral strategies tailored to the structure and stability of different sensory landscapes^1^. In many instances, animals can effectively navigate using a simple chain of sensorimotor reflexes in which they steer towards or away from a salient object based solely on immediate sensory cues^2,3^. However, when sensory signals are intermittent or unreliable, memory mechanisms may facilitate navigation by allowing animals to store the position of stable landmarks in their environment and use an allocentric representation of space to track towards objects that cannot be continuously sensed. Insects, for example, are thought to use a vector-based navigational strategy in which they integrate directional and distance information to guide them toward a remembered location^3,4^. Such an approach is exemplified by the navigational feats of central place foragers, like ants and bees, who can wander far from their nest and efficiently return even in a largely featureless environment^3,5^. However, whether similar vector-based strategies can be used to steer through a more dynamic sensory environment, like an odor plume wafting from its source, has remained unclear^2,6,7^.

Odors are among the most salient signals that insects use to guide navigation^8^. Plume tracking in insects has been proposed to rely on a largely reflexive strategy in which animals surge upwind upon encountering an odor and cast crosswind upon losing contact with the plume in order to re-encounter it^9–11^. Yet, the intrinsically dynamic nature of olfactory plumes often requires animals to navigate using only brief intermittent odor encounters^2,12^, separated by vast stretches of clean air that that offer no immediate sensory information that could point them back to the plume. Memory mechanisms could allow animals to bridge across their fragmented experience of the odor, for example by enabling them to store the direction of their last odor encounter to guide their returns to the plume. We therefore asked whether *Drosophila* can track a plume using components of the same vector-based navigational strategy that central place foragers use to home to their nests. By developing a virtual reality paradigm for plume navigation, we show that *Drosophila* indeed store the heading direction required to return to the plume’s boundary as a dynamic landmark, continuously updating this goal direction as they track along the plume’s border. Consistent with the notion that a plume is not associated with a fixed point in space, flies appear to retain a memory only indicating the direction to the plume’s boundary, not their distance from it. Our results suggest how elements of a conserved navigational toolkit^13^ can be flexibly deployed in different sensory contexts, allowing flies to use odors as remembered chemical signposts to track long distances towards a remote source.

## Flies track a plume along its boundary

One fundamental challenge to elucidating the behavioral algorithms underlying plume navigation is that odors are invisible and often carried along turbulent fluid flow^2,14^. Consequently, few experimental systems offer access to an animal’s true sensory experience as it navigates an odor plume, or to the underlying neural dynamics used to guide behavior. To overcome this limitation, we developed a closed-loop olfactory paradigm that allows head-fixed *Drosophila* to navigate virtual chemical landscapes. In this system, the heading of a tethered fly walking on an air-supported ball is yoked to the rotation of a tube carrying a constant airstream, enabling the fly to control its orientation relative to a wind source blowing from a constant direction in an external or allocentric reference frame^15,16^ (**Fig. 1a, Supplementary Video 1**). By coupling two high-speed mass flow controllers to the virtual XY position of the fly, we could dynamically control the concentration of odor infused into the airstream at each moment as the fly explored this two-dimensional virtual world, allowing us to construct plumes of almost any spatial structure. Notably, in this paradigm flies walk in complete darkness and thus can only navigate using spatial information conveyed by the direction of the wind source and, when they are in the plume, the identity and concentration of the odor. This virtual paradigm thus offers precise control of a fly’s sensory environment, allowing us to investigate how navigational strategies are shaped by ongoing or past odor encounters.

**Figure 1:**
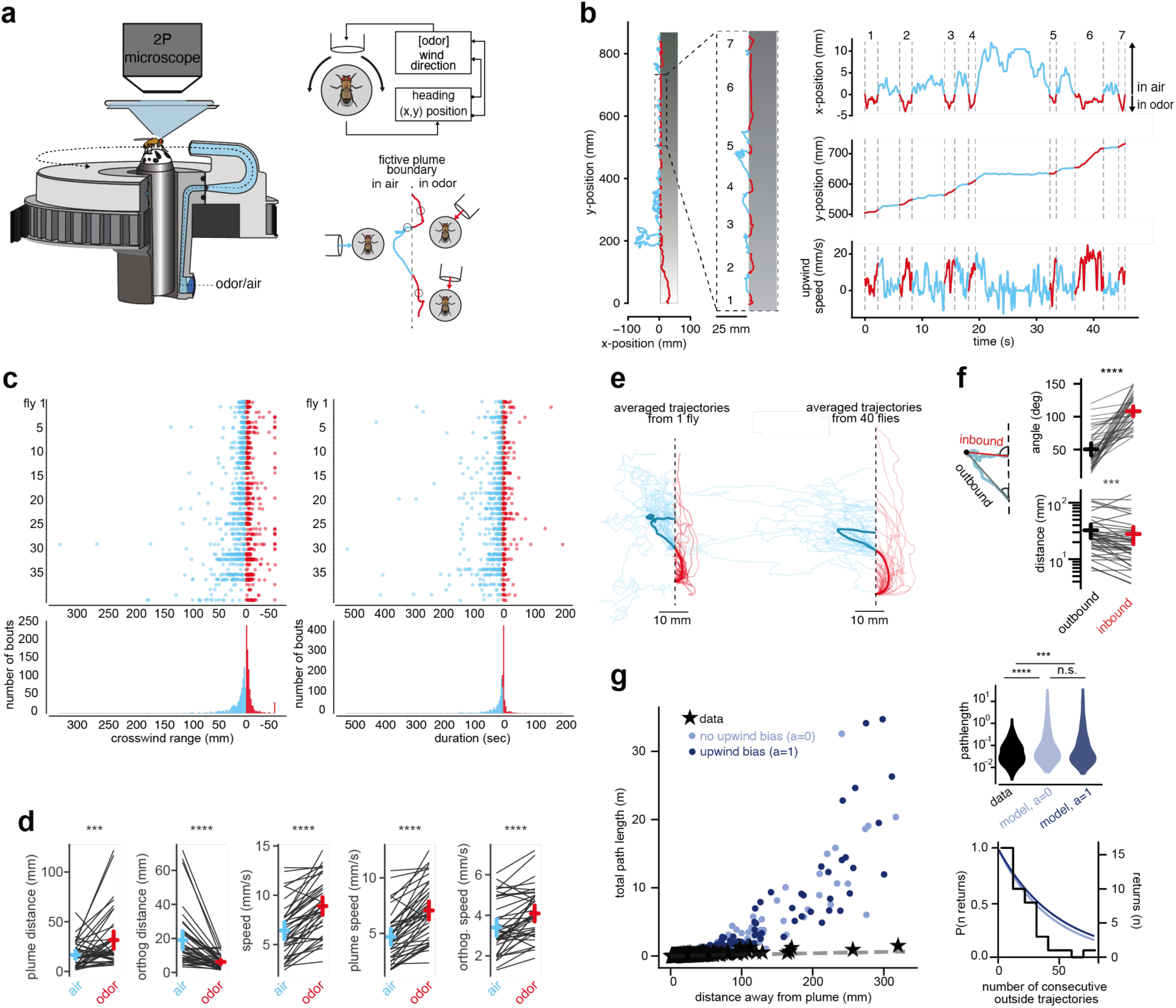
A virtual reality system for olfactory navigation. **(a)** Left: Schematic of the closed loop virtual reality system for plume navigation in which a tethered, walking fly’s heading on an air-supported ball is used to control the angular position of a nozzle delivering a constant airstream, with or without odor. Right, above: The fly’s fictive position is used to control the odor concentration delivered through the airstream. Right, below: Schematic depicting how the position of the airstream and odor concentration change as a head-fixed fly navigates across the fictive plume boundary (dashed line). Trajectory is red when the fly is in the odor and blue outside the odor corridor. **(b)** Left: an example trajectory of a fly tracking a 50 mm wide odor corridor with an ascending gradient (10-100% apple cider vinegar over 1 m). Trajectory is red when the fly is in the odor and blue outside the odor corridor, with enlarged view of the highlighted segment delineated by dashed box. Right: x-position (crosswind axis), y-position (upwind axis), and upwind speed for the inside/outside bouts shown in the highlighted segment. **(c)** Distribution of crosswind distances (left) and durations (right) for the inside and outside bouts of 40 flies. Each dot represents a single bout inside the odor corridor (red) or outside the odor in clean air (blue). **(d)** Comparison of average behavioral metrics for all bouts inside the odor (red) or outside the odor in clean air (blue). Plume distance refers to distance traveled along the longitudinal plume axis while orthogonal distance refers to distance traveled orthogonal to the plume. Each line represents the average for all inside or outside bouts for an individual fly, with the average across flies ±S.E.M. shown in red or blue. n=40 flies. **(e)** Aligned trajectories inside (red) and outside (blue) the plume. Left: all trajectories (thin lines) for the fly shown in (b) and the scaled average for that individual (thick line). Right: Aligned scaled average trajectories for 40 flies (thin lines) and average across all animals (thick line). **(f)** Comparison of the inbound and outbound segments of outside trajectories, defined as leading away from (outbound, black) or returning to (inbound, red) the point farthest from the plume in the crosswind direction (see Methods). Top right: Angles between outbound and inbound segments and plume as depicted in schematic at left. Bottom: Pathlength of outbound and inbound segments, plotted on logarithmic scale. Each thin line represents the average for all trajectories of an individual fly, with the inter-individual average ±S.E.M. shown. n=40 flies. **(g)** The total pathlength of an outside bout as a function of crosswind distance away from the plume for fly data (black stars) or a random-walk model (see Methods and Extended Data Fig. 5) with (dark blue) and without an upwind bias (light blue). Dashed line represents maximal possible efficiency (e.g. when the pathlength is twice the crosswind distance indicating a straight path away from the plume and back through the shortest pathlength). Right top: the distribution of path lengths from real flies (black) and random models, without an upwind bias (dark blue) and with upwind bias (light blue). Right bottom: the probability of returning for *n* consecutive returns with data from real flies (black) and random models, without an upwind bias (dark blue) and with upwind bias (light blue). n=755 outside trajectories from 40 flies and 755 simulated trajectories. n.s. not significant; ***p < 0.001; ****p < 0.0001. Details of statistical analyses and sample sizes are given in Table S1.

We reasoned that memory mechanisms to facilitate navigation might be most apparent in the context of a stationary plume which, through its stability, would allow flies to accumulate useful information about its structure. We therefore initially explored how flies navigate a 50 mm fictive ‘corridor’ of the appetitive food odor apple cider vinegar oriented parallel to the wind direction (ACV, **Fig. 1b**)—a highly simplified chemical landscape that nevertheless recapitulates several key features of the slowly dispersing surface plumes that walking flies encounter in nature when close to their source^12^. The odor corridor was bounded on both sides, such that if a fly walked laterally (crosswind) too far in one direction or the other, it would exit the boundaries of the fictive plume and the ACV concentration would decrease rapidly (**Fig. 1a,b**)). Due to the finite time required for odor entering the airstream to reach the fly’s antenna (∼500 ms), the lateral boundaries of the fictive plume were not perfectly sharp but decayed within ∼5 mm, depending on the walking speed of the fly (**Extended Data** Fig. 1b). A shallower gradient ran along the longitudinal direction of the corridor, such that a fly would experience a steadily increasing concentration of ACV (10-100% over 1 meter) as it walked upwind, mimicking the sensory feedback of getting closer to the odor source.

Prior descriptions of plume tracking suggest that flies may navigate this corridor through odor-gated anemotaxis, reorienting and surging upwind upon contacting the odor corridor, while remaining largely confined within its boundaries^1,9–11,17,18^. However, we found that tethered flies reliably ascended the entire length of the plume by tracking along a single edge through the repeated implementation of two behavioral motifs: upon entering the corridor, they rapidly counter-turned to exit it, and once outside the plume they performed a more circuitous exploration before re-entering (**Fig. 1b, Supplementary Video 2**). We call this behavior ‘edge tracking’.

During edge tracking, flies traveled inside the plume briefly, spending only around 2.01 sec (mean, 95% confidence interval: 1.91 to 2.15 sec, n=755 bouts) in odor before exiting the plume’s boundaries (**Fig. 1c**). As a consequence, animals spent significantly more time outside the odor corridor than inside of it as they ascended upwind along the plume’s boundary. Nevertheless, most of their progress towards the fictive odor source occurred during these brief inside bouts (**Fig. 1b,d**) as flies accelerated upwind upon entering the odor, prior to counter-turning and leaving it. On average, flies also progressed upwind when they walked outside the plume (**Fig. 1b,d**), but these outside trajectories were far more variable in both distance and duration, with some excursions extending hundreds of millimeters orthogonal to the plume’s boundary before returning (**Fig. 1c,d**). Flies typically tracked along a single edge, spending most of their time within the lateral 4.65 mm margin (mean, 95% confidence interval, 4.54 to 5.14 mm) of the odor corridor and rarely crossing the plume’s width to the other edge. Indeed, >96% of the time flies entered the plume, they exited along the same edge (n=793 inside bouts, n= 40 flies). The majority of flies initially encountering an ACV corridor spontaneously tracked its edge for hundreds of millimeters (**Extended Data** Fig. 2), underscoring that edge tracking represents a robust strategy to navigate spatially structured chemical landscapes.

Animals can use perceived changes in odor concentration, sampled either spatially or over time, to localize an odor source^1^. To assess whether edge-tracking flies relied on the increasing odor concentration experienced as they ascended the plume, we provided them with a corridor of the same dimensions but with a constant odor concentration (20% ACV). Individual flies tracked this constant concentration corridor as effectively as plumes in which a longitudinal gradient was present (**Extended Data** Fig. 3a,b), indicating that feedback about increasing odor intensity was not necessary to sustain continued pursuit towards the fictive odor source. Indeed, flies even tracked the edge of a corridor in which the odor gradient was reversed such that as animals progressed upwind, they encountered successively lower concentrations of ACV (**Extended Data** Fig. 3a), a configuration rarely encountered in nature. On average, the trajectories of flies were indistinguishable whether they tracked plumes of constant, increasing or decreasing odor concentration (**Extended Data** Fig. 3b). Flies in our paradigm thus appear sensitive to the presence of odor but not its instantaneous concentration. These observations align with the notion that in naturalistic plumes the intermittent packets of odor that a fly may encounter rarely contain concentration gradients that point to the odor source or can be used to guide navigation^14,19,20^. Flies in our paradigm thus do not appear to rely on the gradual changes in odor concentration along the plume’s longitudinal axis to maintain pursuit. Rather the lateral boundaries of the plume, the region of highest contrast between odor and no odor, may represent a key sensory feature that guides plume tracking.

Edge tracking is reminiscent of descriptions of insects and other arthropods that track the boundary of a surface plume using bilateral comparisons of their chemosensory appendages^21–27^. However, the boundaries of our fictive plume arise from rapid temporal changes in odor concentration that are nearly synchronous between the left and right antenna (**Extended Data** Fig. 4a). To assess whether flies use transient asymmetries in the sensory signals to their two antennae, we expressed the optogenetic activator CsChrimson in the majority of olfactory sensory neurons using the *Orco* promoter, allowing for simultaneous activation of both antennae as flies weaved in and out of a 660 nm light corridor (**Extended Data** Fig. 4b). In the absence of wind, Orco>CsChrimson flies preferentially remained within the light corridor (**Extended Data** Fig. 4c,d), consistent with the intrinsically appetitive nature of broad olfactory sensory neuron activation^22,28,29^. However, these flies progressed minimally in either direction along the plume’s longitudinal axis (**Extended Data** Fig. 4e), further underscoring that the wind provides a critical directional cue to propel animals along the boundary. Indeed, when we added a closed-loop wind source, Orco>CsChrimson flies ascended upwind, adhering to the boundary of the fictive odor plume using a similar behavioral strategy as observed during edge tracking of an ACV corridor (**Extended Data** Fig. 4c,d). Flies thus do not appear to require a bilateral comparison of olfactory input to localize the plume’s edge but rather rely on wind direction and the steep change in odor concentration at the lateral boundary as spatial cues to edge track.

## Edge tracking is not a random search process

In our setup, once a fly is outside the plume, the wind is the only sensory signal that can be used as a directional reference, which in our virtual paradigm blows from a constant allocentric direction. While many flies in our experiments closely adhered to the plume’s edge, with only transient outside excursions, some occasionally performed long trajectories that placed them hundreds of millimeters away from the plume’s boundary (**Fig. 1c**). During these distant excursions, flies were immersed in clean air for tens of seconds before returning back to the edge, frequently through a shorter more-directed path (**Fig. 1c-e**), suggesting that they may retain information about the direction or location of the plume to guide their returns. To examine the structure of a fly’s exploration outside the odor corridor, we divided each outside trajectory into an ‘outbound’ and ‘inbound’ segment based on the most distant point from the plume’s boundary. Flies typically exited and returned to the plume’s edge using distinct routes. During outbound trajectories, flies left the plume at a shallow angle relative to the plume’s edge, steering strongly upwind. In contrast, inbound trajectories were often angled almost perpendicular to the plume’s edge and followed shorter paths such that, on average, flies took more direct paths back to the odor corridor (**Fig. 1f**). Calculating the average path of all the trajectories for an individual fly further supported the asymmetry of their routes away from and back to the plume’s edge (**Fig. 1e**).

The observation that animals take lengthy excursions away from the plume and make directed returns back to its boundary suggests that their outside exploration may not be random. To gain further insight into the structure of a fly’s trajectories outside the plume, we decomposed them into their elementary run lengths and turn angles (**Extended Data** Fig. 5a,b Methods), then randomly drew from these experimental distributions to generate synthetic trajectories built from the same path statistics but lacking any directional biases, analogous to the reflexive and random casting that insects have been proposed to perform when they lose contact with a plume^1,30^. Simulated trajectories could reliably return to the plume’s boundary (**Fig. 1g**) but displayed several differences from the trajectories of real edge-tracking flies. Notably, while simulated trajectories could extend far away from the plume, they were highly circuitous and took significantly longer overall path lengths to return. For example, in our dataset, flies walked up to 320 mm from the lateral boundary of the plume, taking a total roundtrip path of 1.5 m—a distance just over twice the length of the most efficient path they could achieve by walking directly away from the plume and directly back. Simulations of similarly lengthy outside excursions, by contrast, took ∼15 times longer than the most efficient paths (**Fig. 1g**). Additionally, while the outside trajectories of real flies were significantly shorter on the inbound than outbound segments, reflecting more directed returns, the outbound and inbound segments of simulated trajectories were equivalent in length, consistent with their undirected nature (**Extended Data** Fig. 5d). Finally, real flies display an intrinsic upwind bias^16,31,32^ and continued to walk upwind even outside the plume. Adding the experimentally observed upwind bias (**Extended Data** Fig. 5c) resulted in simulated trajectories that ascended the plume like those of real flies. However, this upwind bias had no impact on the efficiency of simulated trajectories (**Fig. 1g, Extended Data** Fig. 5c,e), consistent with the fact that the most efficient path to return to a vertically-oriented odor corridor depends only on the ability to progress in the crosswind, not upwind, direction. Edge tracking therefore does not adhere to the statistics of a random search process. Rather, the directed returns to the plume’s edge observed in edge-tracking flies suggest they rely on a memory of past plume encounters to guide efficient trajectories back to the boundary.

## Flies can track plumes with different geometries

The nature of memory mechanisms that facilitate navigation back to the plume could take several forms, from simply biasing the direction of an animal’s search, as observed after a hungry fly consumes a sugar droplet^33,34^, to storing the specific location of the plume’s boundary and homing towards it via path integration, as displayed in foraging ants returning to their nest^5,35^. To shed light on the algorithms that flies may use for edge tracking, we leveraged the virtual nature of our paradigm to vary the structure of the fictive chemical landscape, by changing either the orientation and stability of the plume or modifying the fly’s distance to it.

Transient shifts in wind direction may cause a plume to meander, such that it no longer aligns with the wind direction or points to the odor source. Employing a reflexive behavioral strategy in which an animal surges upwind upon encountering the odor would inevitably cause it to walk out of a meandering plume, leading to the proposal that it may be advantageous to track the envelope of the plume rather than the wind direction^36^. To explore this possibility, we constructed a set of ‘tilted’ plumes of constant ACV concentration, in which the longitudinal axis of the 50 mm corridor was offset from the wind direction by either 45 or 90 degrees (**Fig. 2a-c**). We found that flies robustly tracked the boundaries of these tilted plumes for many hundreds of millimeters, even for the 90-degree plume which required flies to initially choose a left or right crosswind direction and progress along it, moving orthogonal to the wind with minimal backtracking. Indeed, flies even tracked the edge of a 90-degree plume along a diminishing concentration gradient (**Extended Data** Fig. 3c,d), reinforcing that their progression does not depend on the instantaneous odor concentration. Rather flies appear to use an internal sense of direction allowing them to advance in a given direction for hundreds of millimeters.

**Figure 2:**
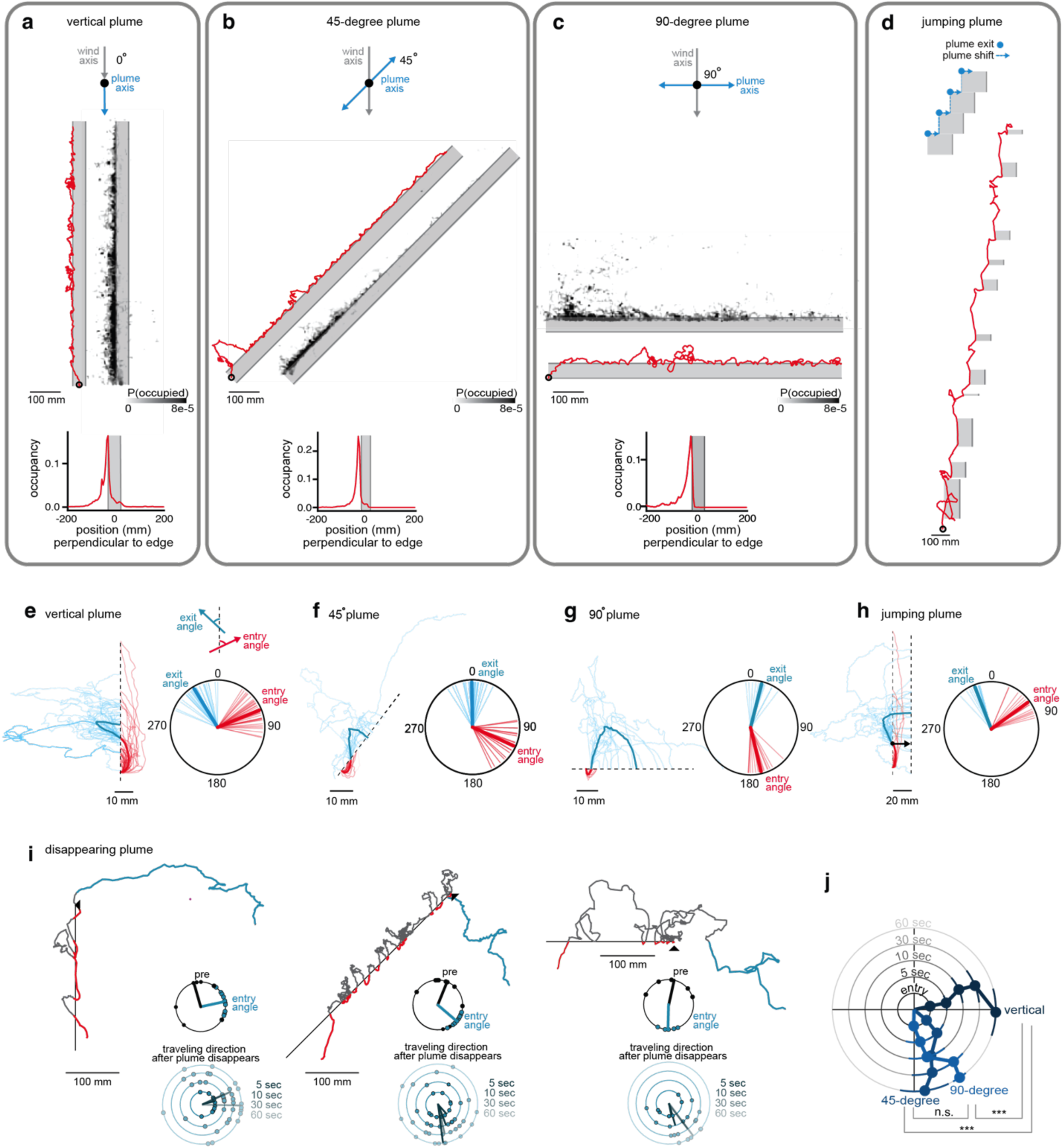
Flies track plumes that are not aligned with the wind. **(a-c)** Flies track plumes with different orientations relative to the wind. For each plume type, we show a single representative trajectory (red, with starting position denoted by black circle); heatmap in greyscale depicting occupancy, and occupancy as a function of distance perpendicular to the plume. **(a)** n=40 flies; **(b)** n=21 flies; **(c)** n=20 flies. (**d**) Representative trajectory of a fly navigating a ‘jumping’ plume where the plume shifts 20 mm to the right, each time the fly exits the plume. Trajectory is shown in red with starting position denoted by black circle. **(e)-(h)** Left: aligned and averaged inside and outside trajectories for each plume orientation. Each thin line represents the scaled average for a single individual and the thick line represents the average across all individuals. Right: distribution of entry and exit angles for each plume type. Each thin line represents the average for all exits and entries from a single fly and the thick line represents average across all individuals. **(e)** n=40 flies; **(f)** n=21 flies; **(g)** n=20 flies; **(h)** n=24 flies. Entry and exit angles were determined by calculating the mean heading in the 0.5 sec prior to and 0.5 sec after crossing the plume’s boundary as shown in (e). **(i)** Trajectories of flies after the plume vanishes while the fly is on outside bout the plume for 0-degree (vertical), 45-degree and 90-degree plume. For each plume orientation, a representative example is shown with the plume’s boundary indicated by black line, and inside bouts shown in red and outside bouts in grey. The point at which the plume disappears is indicated by the black arrowhead, with the fly’s trajectory over the next two minutes shown in blue. For each, top polar plot shows the distributions of average heading direction while animals walked in closed loop wind in the absence of odor during the two minutes prior to plume tracking (pre, black) and average entry angle during plume tracking (blue). Bottom polar plot shows the average heading direction at indicated times after the plume’s disappearance. Each dot represents an individual fly with vertical n=12 flies; 45-degree n=11 flies; 90-degree n=6 flies. **(j)** Comparison of the average entry angle during plume tracking (innermost circle) and traveling direction for the indicated times after the plume’s disappearance for all flies in (i). Each circle reflects the indicated time bin, with mean and SEM shown. Details of statistical analyses and sample sizes are given in Table S1. n.s., not significant, ***p < 0.001. Details of statistical analyses and sample sizes are given in Table S1.

Flies tracked the boundaries of both 45- and 90-degree tilted plumes along their upwind edge using a similar behavioral strategy as the vertically oriented plume (0-degree): upon entering the odor corridor, they rapidly reoriented to steer upwind, and upon losing contact with the plume, they performed directed returns back to the plume’s boundary (**Fig. 2e-g, Extended Data** Fig. 6a,c**-e**). However, flies adopted distinct angles to enter and exit a plume that depended on its geometry. For example, to exit a vertical odor corridor while maintaining an upwind bias requires that animals steer laterally away from the upwind direction to counterturn out. By contrast, flies could leave the 45- and 90-degree plume using a large range of upwind angles, including directly upwind. Yet, on average individuals did not use the full range of possible exit angles that could maintain an upwind bias, but rather adopted a relatively narrow range of exit angles that were frequently biased in the direction that they were progressing along the plume’s length (**Fig. 2e-g, Extended Data** Fig. 6a,c**-e**). Likewise, entry angles also depended on the plume’s orientation, reflecting the fact that returning to a vertical plume requires animals to walk in a crosswind direction, but returning to a 90° plume requires them to direct their trajectories downwind. Inside trajectories were shorter in tilted plumes compared to vertical plumes (**Extended Data** Fig. 6b), as flies exited more rapidly when steering upwind within the odor. In contrast, trajectories outside of tilted plumes were on average the same length even when flies had to counter their intrinsic upwind drive^16,31,32^ outside the plume to return back to its boundary **(Extended Data** Fig. 6b).

Comparing simulated trajectories—generated by randomly selecting from the experimental distribution of run lengths and turn angles—with the paths of real flies underscored the importance of these biases to effectively track a tilted plume (**Extended Data** Fig. 5f-g). As observed for a vertical plume, simulated trajectories that lacked an upwind bias were frequently able to return to the boundary of a 45- or 90-degree plume, although far less efficiently than real trajectories. However, incorporating the inherent upwind bias displayed by real flies as they walk outside the plume (**Fig. 1b,d**) resulted in simulated trajectories that strayed far away from the boundary and often failed to return (**Extended Data** Fig. 5f-g), emphasizing that successfully tracking a tilted plume requires flies to counter their upwind drive to reenter. Flies thus appear to adapt their trajectories to the geometry of the plume. Even in clean air, where flies have nothing but the wind direction to serve as a navigational cue, they adopt different biases to return to the plume depending on its geometry—directing their trajectories crosswind to return to a 0-degree plume but steering downwind to return to a 90-degree plume (**Fig. 2e-g, Extended Data** Fig. 6c**-e)**.

To determine whether flies rely on a memory of the direction of the plume’s boundary to guide their returns, we allowed flies to track the edge of a vertical, 45- or 90-degree plume for 10 minutes before removing the plume while they were on an outside excursion (**Fig. 2i**). We found that after the plume’s disappearance, flies continued to walk in the same direction that they had previously used to return to the plume and maintained this distinct directional bias for tens of seconds, even though this often took them hundreds of millimeters away from the location of their last plume encounter. The directional biases differed across the three plume geometries and were distinct from the upwind bias that flies generally displayed in the two minutes prior to encountering the plume (**Fig. 2i,j**), even though they had continuous access to the same closed-loop wind source to orient them. Thus flies appear to form and maintain a persistent directional bias that depends on the orientation of the plume that they had been previously tracking. While the disappearance of a sugar droplet has been shown to trigger flies to perform a local search centered around the previously encountered reward location^34^, flies did not dwell around the location of their last plume encounters, suggesting that edge tracking does not involve positional information.

One implication of storing a directional bias back to the edge is that it should allow flies to effectively track dynamic plumes, even as they shift over time. To explore this possibility, we jumped the position of the plume each time a fly exited it by moving the boundary 20 mm in the direction opposite to their exit (**Fig. 2d,h**). Successful navigation of this dynamic plume requires flies to not only bias their return trajectory in the direction of the plume’s boundary, but to walk even further in this direction to re-encounter it, resulting in significantly longer trajectories than for stationary plumes (**Extended Data** Fig. 6b). Flies readily returned to and tracked the jumping plume (**Extended Data** Fig. 5h**, Supplementary Video 3**), consistent with the persistent directional bias they displayed when the plume disappeared, allowing them to return to a plume even as it meandered away from the last encountered position. Notably, flies did not search for the plume boundary at its pre-shifted location, providing further support for directional but not positional memory.

## Edge tracking requires the fly compass system

Our behavioral analyses suggest that flies rely on a directional memory of the plume’s boundary to bias their outside trajectories and direct their returns. We reasoned that the central complex, an array of evolutionarily conserved neuropils underlying spatial navigation in insects^3^, might serve as an ideal neural substrate for encoding and storing memories of the plume’s direction that could be used to guide edge tracking. A core spatial signal encoded in the *Drosophila* central complex is a fly’s heading direction, which is represented as a single bolus of activity that rotates around topographically organized EPG neurons in the ellipsoid body^37^. This neural compass can be anchored to stationary visual landmarks, such as the sun^38–40^ or mechanosensory landmarks such as a wind source^41^ to allow animals to maintain a stable heading as they fly or walk in a fixed direction^32,39,42,43^. Edge-tracking flies navigate in complete darkness, leaving the wind direction as the sole external landmark available to orient themselves as they weave in and out of the plume.

Ascending pathways conveying information about wind direction are thought to entrain the activity of EPG neurons^41^, allowing flies to use a stable wind source to maintain an accurate allocentric representation of their heading in the world. To examine the activity of EPG neurons during edge tracking, we performed two-photon functional calcium imaging of their axon terminals within the protocerebral bridge, a linear neuropil comprised of 18 glomeruli that receives two duplicated copies of an animal’s heading direction (**Fig. 3a**). Consistent with prior recordings of EPG neurons during visual navigation tasks, we observed two discrete boluses of neural activity that moved in concert across the left and right halves of the protocerebral bridge, with their position locked to the fly’s heading direction relative to the wind (the EPG ‘phase’). In flies walking in the absence of external visual landmarks that can be used for corrective feedback, the EPG phase can rapidly drift relative to the animal’s true behavioral heading^37,44,45^. However, during edge tracking, the EPG phase remained faithfully aligned to the fly’s heading direction in the virtual environment over the course of tens of minutes (**Fig. 3b,c**), underscoring their ability to use a wind source as a stable landmark^41^. The amplitude and width of the boluses of EPG activity in the protocerebral bridge were comparable throughout the course of an edge-tracking trial, indicating that this heading signal remains yoked to the wind and unchanged by the presence of odor (**Fig. 3d**).

**Figure 3:**
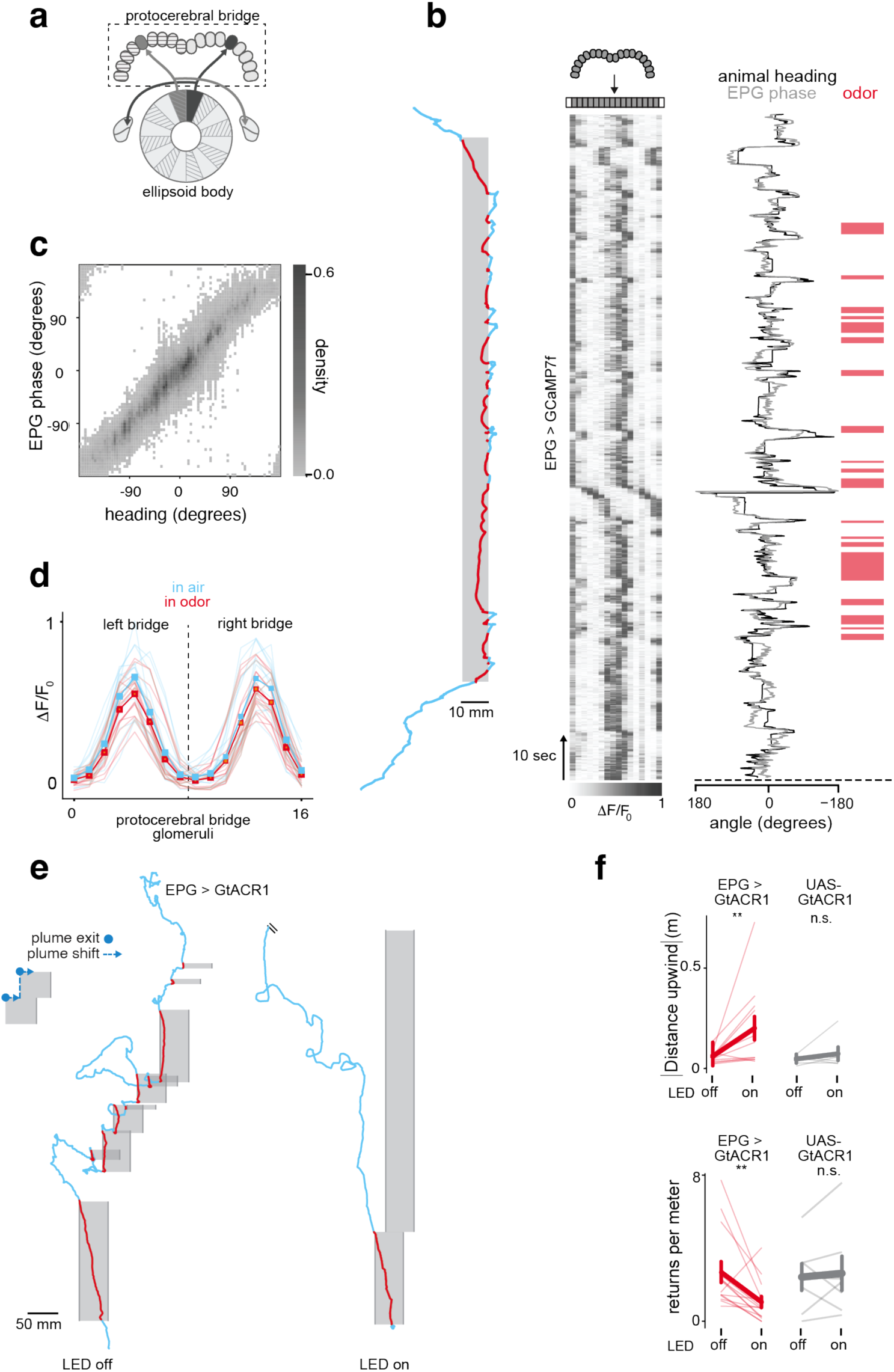
EPG neurons represent a fly’s heading and are required for edge tracking. **(a)** Schematic of the central complex neuropils innervated by EPG neurons. Adjacent EPG neurons in the ellipsoid body project to glomeruli in the left and right protocerebral bridge. **(b)** Representative recording of EPG activity as a fly edge tracks a 10 mm wide vertical plume. Left: trajectory of an individual fly, with inside bouts shown in red and outside bouts in blue. Middle: Heatmap of EPG activity (ΔF/F_0_) during edge tracking. Each column in the heatmap represents EPG activity in 16 glomeruli in the protocerebral bridge as indicated at top. Right: alignment of the EPG phase estimate (grey) with the animal’s heading (black) during this same edge-tracking trial. Timing of in odor bouts are depicted in red at far right. **(c)** Heat-map representing the correspondence between an animal’s heading and the estimated phase of the EPG bump during edge tracking. N=16 trials from 9 animals. **(d)** The amplitude of phase-offset activity bumps in the left and right protocerebral bridge during edge tracking when the animal is walking in the odor (red) or in air (blue). N=16 trials from 9 animals. **(e)** Representative experiment showing the trajectory of an EPG>GtACR fly navigating a jumping plume in the absence (left) and presence (right) of optogenetic inhibition via LED illumination throughout the edge-tracking trial. **(f)** Comparison of distance traveled upwind upon re-encountering the plume (top) and number of successful return trajectories (bottom) for EPG>GtACR and UAS-GtACR control flies in the absence and presence of optogenetic inhibition. Each thin line represents an individual fly, with thick line showing mean across flies and SEM. n.s., not significant, **p < 0.01. Details of statistical analyses and sample sizes are given in Table S1.

The representation of an animal’s current heading by EPG neurons is thought to be integral to *Drosophila* navigation as this signal may be transformed and reformatted to generate other spatial variables, such as an animal’s traveling direction^46,47^ or compared to an animal’s stored goal direction to guide its moment-to-moment steering behavior^32,43,48–50^. To assess whether edge tracking relies on this core navigational signal, we used an intersectional genetic driver to express the light-gated anion channel GtACR1 in EPG neurons and acutely inhibit their activity during edge tracking. We focused on flies tracking a jumping plume given that successful navigation requires animals to maintain a persistent bias to return to the plume’s shifted edge (**Fig. 2d,h**) and may thus reveal the need for a continuous representation of an animal’s heading. Indeed, upon silencing of EPG neurons, flies no longer made directed returns to the plume’s shifted boundary but instead frequently wandered away in the upwind direction (**Fig. 3e**). Consequently, in trials in which EPG neurons were inhibited, flies made fewer returns to the plume’s boundary (**Fig. 3f**). When they did manage to return, they took longer, often entering the plume hundreds of millimeters farther upwind in contrast to the directed, crosswind returns observed in control flies (**Fig. 3e,f**). No differences in the frequency of returns or upwind entry position were apparent in control animals (UAS-GtACR1), independent of whether the optogenetic stimulating LED was on or off, supporting the specificity of this manipulation. EPG neuron activity thus appears to be required for edge tracking. Navigation of structured chemical landscapes therefore appears to represent a spatial navigation task that depends on core elements of the central complex circuitry.

## Flies rely on odor entry and exit angles to guide plume navigation

What spatial cues might flies use to form or update a memory of the plume’s direction during edge tracking? While flies could in theory enter or exit an odor corridor over a 180-degree range, they consistently adopted a narrower range of angles that depended on the plume’s geometry (**Fig. 2e-h, Extended Data** Fig. 6c-f). We therefore hypothesized that each time a fly crossed the edge of the plume, it used the angle of its current heading relative to the wind direction to form an angular memory that could guide subsequent trajectories. If so, odor encounters uncoupled from their corresponding entry or exit angles should lead to different trajectories outside the plume. To investigate this possibility, we allowed a fly to track the boundary of an odor corridor for 10 minutes and then immediately replayed the identical temporal sequence of odor pulses back to the same individual or to a naive animal that had never edge tracked (**Fig. 4a**). During these ‘replay’ epochs, the wind source remained unchanged and yoked to the fly’s angular heading, but the fly’s odor experience was now completely uncoupled from its actions and, in particular, from the angle that the fly was traveling at the time of odor onset or offset.

**Figure 4:**
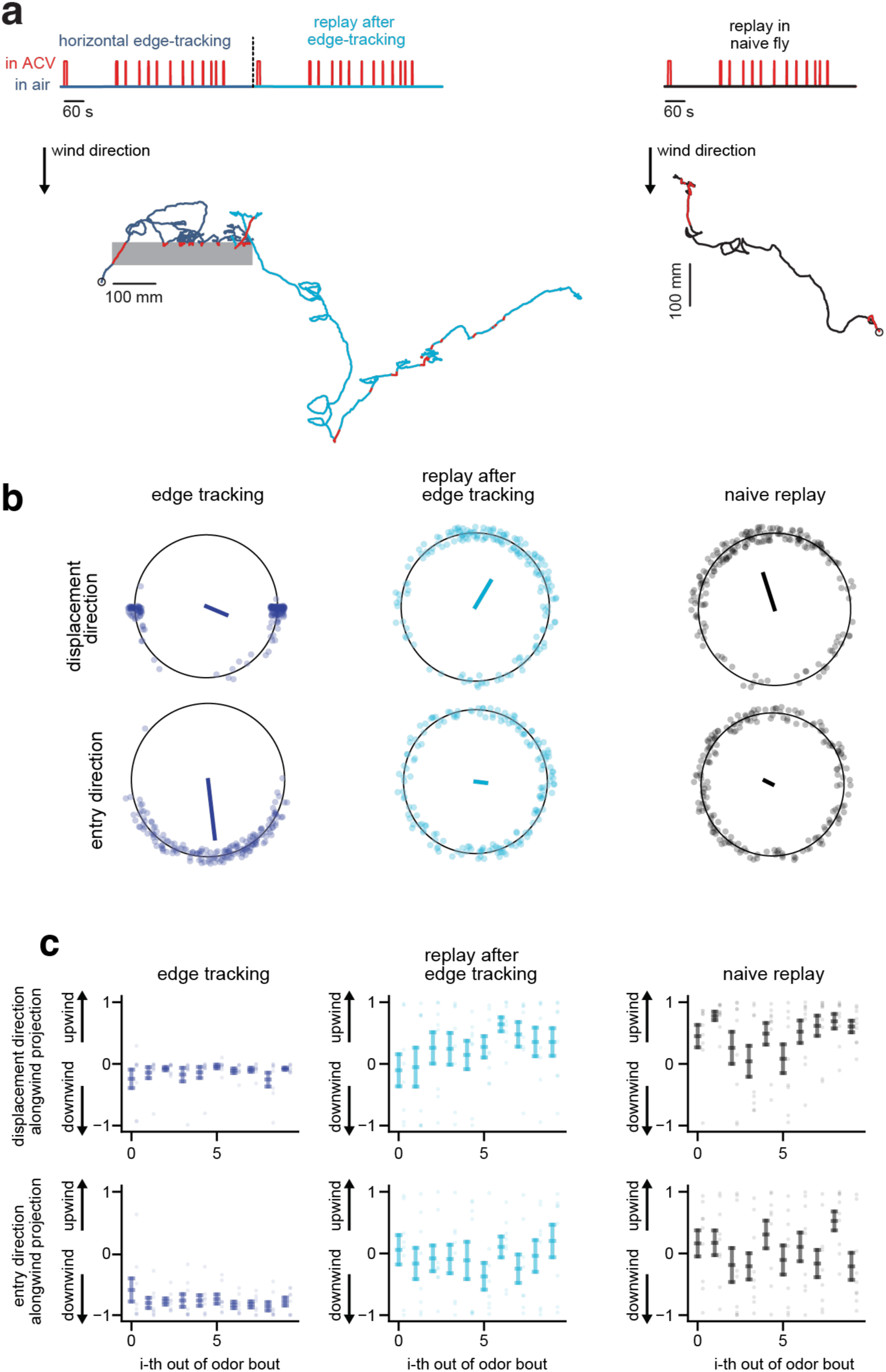
Fly trajectories depend on the spatial structure of their olfactory experience. **(a)** Left: Representative ‘replay’ experiment in which a fly was allowed to edge track along a 90° plume for 10 minutes, after which the same temporal sequence of odors was presented back to the same fly. Schematic on top depicts the sequence of odor encounters during edge tracking that was replayed to the fly during the replay epoch. Representative trajectory during edge tracking shown as red in odor and dark blue when outside the odor, while trajectory during replay epoch is red inside the odor and cyan when outside the odor. Right: Trajectory of a naïve animal when presented with the same odor sequence. **(b)** Polar plots showing the net displacement direction when flies are outside the odor (top,) and entry direction when they encounter the odor (bottom) during edge tracking (left, dark blue), replay to the same fly (middle, cyan), or replay in naïve animals (right, black). Each point represents a single bout for an individual fly, with average vector for all flies shown as a line. n=8 flies. **(c)** The displacement direction and entry direction of flies over sequential outside bouts of edge tracking trials (left, dark blue), replay of the odor sequence to the same flies (middle, cyan), and replay of the odor sequence to a naïve flies (right, black). n=8 flies. The alongwind projection is shown, with 1 denoting straight upwind and -1 denoting straight downwind, to emphasize the change in the direction of outside bouts depending on the conditions in which the fly encounters the odor.

We focused on flies tracking along the boundary of a 90-degree plume given that this requires animals to bias their outside trajectories downwind to reenter the plume, which allowed us to uncouple the biases in an animal’s trajectories arising from its intrinsic upwind drive^16,32^ from those emerging from their recent edge-tracking experience. During the replay epoch, we found that flies that had previously tracked along a 90° plume initially continued to display the same directional biases that they had used to return to the odor corridor: upon odor onset they steered in the upwind direction, while at odor offset they counterturned to travel in a downwind direction (**Fig. 4**). However, this downwind bias rapidly decayed with subsequent encounters, and the traveling direction of flies during replay progressively shifted towards the upwind direction (**Fig. 4c**). In contrast, naive flies presented with the same temporal sequence of odor pulses surged upwind at odor onset and continued to travel in an upwind direction after odor offset, giving rise to trajectories with a net upwind bias (**Fig. 4a**, right). Thus while naive flies track upwind, analogous to prior reflexive descriptions of plume tracking^9,17,18^, individuals that had recently tracked the edge of a 90-degree plume maintained a directional bias based on their past plume encounters that gradually eroded. Together, these results suggest that the paths that flies take as they weave in and out of an odor plume depend on the spatial structure of their previous odor encounters, such that even identical temporal patterns of odor stimulation can give rise to distinct trajectories depending on the fly’s orientation at the time of odor onset or offset. Based on these observations, we conclude that edge tracking flies use their recent entry and exit angles to form and update their goal direction, allowing them to navigate away from and back to the plume’s boundary.

## A behavioral model of edge tracking

To gain further insight into how flies use their past plume encounters to shape their navigational goals, we developed a switching state space model^51,52^ in which simulated animals alternate between ‘leaving’ and ‘returning’ states as they progress along a plume’s boundary (**Fig. 5a**). These states do not correspond directly to the epochs when the fly is inside or outside the plume, although the transition rates between them depend on the presence or absence of odor **(Extended Data** Fig. 7a,b). This model generates a velocity vector for the simulated fly through an auto-regressive process, with a correlation timescale and noise level determined by the current state. Importantly, the velocity vector is biased by a memory of previous exit angles (for the leaving state) or previous entry angles (for the returning state). These memories are updated each time the fly crosses the odor boundary by replacing the current memory with a linear combination of that memory and the direction of motion at the time of the odor crossing. As a consequence, flies in our model rely on entry angles over multiple odor encounters to form and update a goal direction that biases their velocities in the returning state, while they use their exit angles to form and update a goal direction that similarly biases their leaving state. We used variational inference to extract the latent leaving and returning states, and latent goal directions, and to learn the parameters determining the velocity and memory updates from the trajectories of individual flies tracking different plume geometries (**Extended Data** Fig. 7g**, Model Methods**). Model parameters varied across flies, matching the heterogeneity that flies display in their outside trajectories, with some individuals adhering closely to the plume’s boundary and others taking longer and more circuitous excursions before returning (**Extended Data** Fig. 7c-f). From these fits, we computed average model parameters to simulate fly trajectories and reveal how leaving and returning goal directions emerge from repeated plume encounters (**Fig. 5b**).

**Figure 5:**
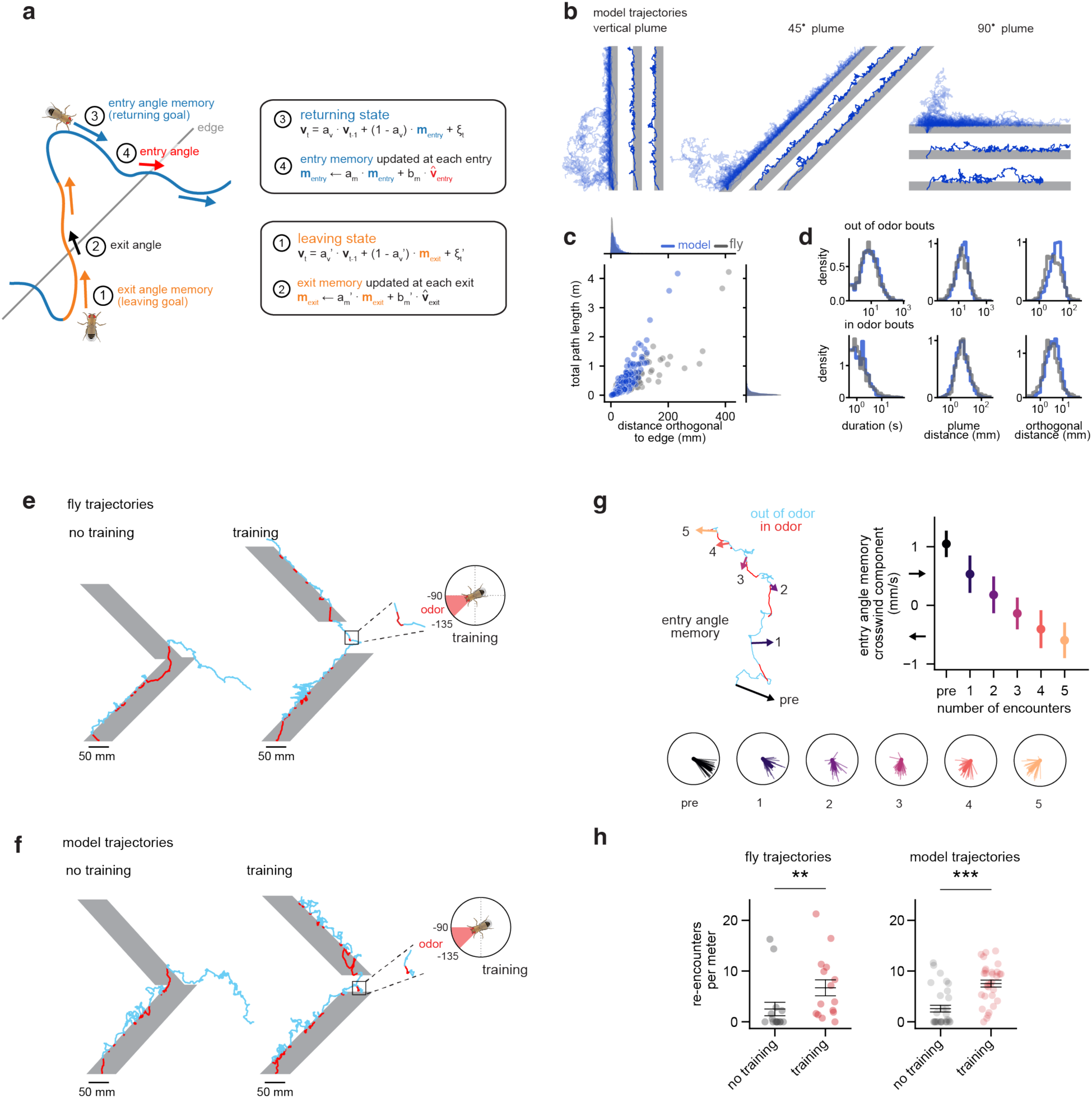
A behavioral model of edge tracking suggests flies rapidly update their entry angle memories with experience. **(a)** Components of the switching space model mapped onto a schematized trajectory of a fly tracking the edge (gray) of a 45-degree plume. When the fly is in the leaving state (orange), it relies on its exit angle memory (m_exit_) which it updates each time it exits the plume (black arrow). When the fly is in the returning state (blue), it relies on its entry angle memory (m_entry_) which it updates each time it enters the plume (red arrow). **(b)** Representative individual trajectories simulated from the model shown for plumes with different geometries relative to the wind. For each plume, two trajectories are shown individually (dark lines) or an overlay of 25 trajectories (semi-transparent lines). **(c)** Comparable efficiency of simulated and real fly trajectories revealed by plotting total path length versus total distance in the direction perpendicular to the plume’s boundary for 72 animals and 75 simulations tracking 0-,45-, and 90-degree plumes. Each dot represents a single outside bout. **(d)** Histograms comparing metrics across population of 72 flies tracking 0-,45-, and 90-degree plumes (gray) or across 75 simulated trajectories (blue). **(e)** Left: Representative trajectory of a naïve fly provided with two 45-degree plume segments demonstrating that in the absence of training (left), flies continue to progress in the direction required to re-enter the first plume segment. Right: A single operant training session (see Methods), in which a fly was exposed to odor when it spontaneously walked in the direction required to enter the second plume segment (-90° to -135°), was sufficient to enable the fly to subsequently track a 45-degree plume segment oriented in the opposite direction. Note that to enter the second plume segment flies must travel in the same direction as used for operant conditioning (-90° to -135°) and so effectively have a second operant training epoch. **(f)** Model simulations of the same experiment shown in (e). **(g)** Top left: Simulation showing how the entry angle memory gradually shifts over multiple odor encounters during operant training paradigm shown in (e), transitioning from pointing to the lower right after tracking the first 45-degree segment to pointing to the lower left. Top right: a plot of the crosswind component of entry angle memory as a function of number of odor encounters during operant training, n=29 simulations, mean +/- std deviation. Bottom: entry angle memory vectors as a function of odor encounters during operant training for same 29 simulations, with individual simulations (thin lines) and average vector (thick line). **(h)** A single operant training trial enhances the ability to track a 45-degree plume segment oriented in the other direction. Shown is the number of plume re-encounters on the second segment 45-degree plume of both real (left) and model (right) flies. **p < 0.01, ***p < 0.001. Details of statistical analyses and sample sizes are given in Table S1.

The model requires that we initialize the memory vectors at the beginning of a simulation. In the absence of information about the prior olfactory experience of a given fly, we set the initial exit memory to be upwind with a narrow randomly chosen crosswind component (see Model Methods), in accord with the observed upwind bias that flies display as they exit all plumes (**Fig. 2e-h, Extended Data** Fig. 6c-f). The memory of the entry angle, on the other hand, was initialized as a null vector, consistent with the assumption that flies have no prior information about which direction to return before encountering a plume. Simulations based on this model capture the ability of flies to track the edges of plumes with different geometries and match the statistics of their experimental trajectories (**Fig. 5b-d**, **Extended Data** Fig. 8). Simulated trajectories from a model lacking an entry angle memory were significantly less effective in advancing along the plume’s boundary for all plume orientations (**Extended Data** Fig. 9a,e). Likewise, simulations using an exit memory that was either purely upwind or contained a crosswind component that was resampled with each exit were also impaired in some plume geometries (**Extended Data** Fig. 9c**-e**). In contrast, a model in which the exit angle memory maintained a constant crosswind bias could edge track successfully (**Extended Data** Fig. 9b,e). These results align with evidence that flies strongly bias their entry angles to track plumes of different orientations (**Fig. 2e-h, Extended Data** Fig. 6c-f). By contrast, flies exhibit less variation in their exit angles, which display a small consistent crosswind bias that is only minimally influenced by a fly’s subsequent edge-tracking experience (**Extended Data** Fig. 9f-g).

Our model provides insight about the timescales of memory formation and updating that shape edge tracking behavior. For instance, the model posits that flies have no initial entry angle memory, but a goal is established and reinforced with repeated plume encounters as an animal repeatedly enters the odor corridor at consistent angles. Indeed, both real and simulated outside trajectories become more efficient over time (**Extended Data** Fig. 10a). This effect was only significant for the 90° and jumping plumes, likely because these configurations cause flies to be displaced further from the plume’s edge. Once an exit or entry angle memory is formed, the model predicts that it will be retained in the absence of additional experience, in accord with our observation that animals continue to travel for tens of seconds in the direction of a previously encountered plume that has suddenly vanished (**Fig. 2i**). On the other hand, simulation of the olfactory replay experiments suggest that an animal’s entry angle memory can be rapidly updated through a small number of additional odor encounters (**Fig. 4d, Extended Data** Fig. 9f**, Extended Data** Fig. 10b,c), allowing animals to flexibly adapt their navigational goals.

To explore the timescales of memory updating, we provided flies with a plume consisting of two 45-degree segments oriented in opposite directions relative to the wind (±45°) and examined how a given entry angle memory adapts due to additional odor encounters in a different direction (**Fig. 5e, Extended Data** Fig. 11a). Successfully tracking both segments requires distinct entry angle memories that are approximately 90 degrees apart. After tracking the first (+45°) plume segment, the majority of flies continued to travel away from the plume, replicating the persistent directional bias observed when the plume disappeared (**Fig. 2i, Extended Data** Fig. 11c). As a consequence, flies rarely re-encountered the second (-45°) segment or succeeded in tracking upwind along its boundary (**Fig. 5h, Extended Data** Fig. 11b,d), consistent with the notion that they require additional olfactory experience to update their goal direction.

We therefore introduced an operant ‘training’ session after animals tracked the first +45° plume segment to update the entry angle memory by exposing a fly to odor when it spontaneously walked in the direction required to enter the second segment’s boundary (e.g. -90° to -135°, **Fig. 5e**, **Extended Data** Fig. 11a). Model simulations indicated that the entry angle memory could be shifted to a new angle by a small number of successive odor encounters at that angle (**Fig. 5g, Extended Data** Fig. 9g). Indeed, flies trained with a single operant session were significantly more successful in tracking the second plume segment than flies without this training, suggesting they incorporated a modified entry angle memory (**Fig. 5f**). Additional operant training sessions in the same direction (5 sequential bouts) provided no further enhancement, whereas flies trained in the opposite direction, by exposing them to odor if they walked in the direction that matched their reentry into the first plume segment (+90° to +135°), performed similarly to untrained flies (**Extended Data** Fig. 11b,d). Together, these experiments support the idea that flies bias their return to the plume’s boundary using an angular memory that can be updated dynamically as the orientation of the plume shifts, allowing for flexible navigation in a shifting chemical environment.

## Dynamic goal signals within the central complex

Our behavioral model has two dynamic components: stochastic transitions between leaving and returning states that depend on the presence or absence of odor, and state-dependent updates of velocity and memory vectors (**Extended Data** Fig. 6a). While we do not attempt to assign the state-determining component of the model to any specific circuitry of the fly brain, its velocity and memory vector representations map well onto the circuitry of the fan-shaped body in the *Drosophila* central complex (**Fig. 6a**). In particular, the fan-shaped body is thought to encode allocentric goal signals that could guide olfactory navigation through its grid-like organization of intersecting columnar and tangential neurons. Columnar neurons of the fan-shaped body integrate wind and heading signals to represent spatial variables^46,47,53,54^, such as a fly’s traveling direction, as vectors through sinusoidal population activity. Tangential neurons receive input from olfactory pathways^49,55,56^ and are thus poised to modulate the vector representations of columnar neurons during plume tracking. Based on this suggestive circuit architecture, we hypothesize that the memory of a fly’s entry direction, as it weaves into and out of the plume, is stored as a sinusoidal pattern in the strengths of plastic synapses between tangential and columnar neurons, analogous to other anatomically-inspired models of angular goal formation in *Drosophila*^48,50^. Memory updates could be driven by tangential neurons that are activated at the time of plume boundary crossings^55^. Distinct odor-responsive tangential neurons could gate the use of these exit and entry angle memories, allowing animals to recall different goal directions when they are in the leaving and returning states. The resulting goal direction is then compared to the fly’s current heading to generate corrective steering signals that guide the moment-to-moment motor patterns of navigation^32,43,48,49^.

**Figure 6:**
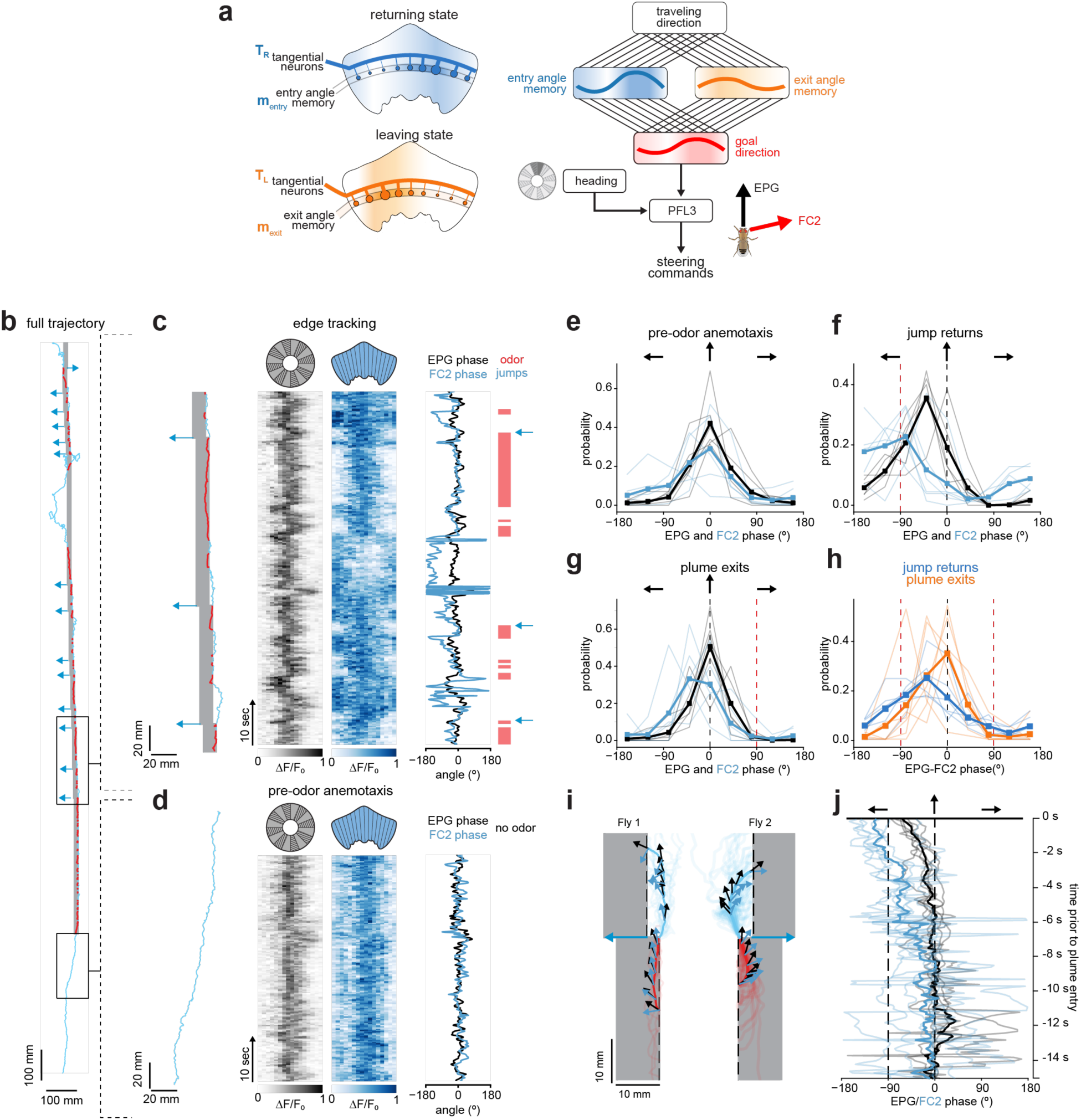
FC2 neurons signal the direction of the plume’s boundary prior to returns. **(a)** Left: Schematic of circuitry in the fan-shaped body hypothesized to store entry and exit angle memories. Memories are stored in synapses between state-activated tangential neurons and columnar neurons encoding traveling direction. Right: Schematic of the full model. Crossing the plume’s boundary drives the storage of an entry or exit angle memory that is alternately read out in the returning or leaving states. Memory is conveyed to FC2 neurons representing the fly’s goal direction, and this direction is compared to the fly’s current heading by the PFL neurons that drive corrective steering to align the fly’s current heading with its goal. **(b)** Representative experiment in which the activity of FC2 neurons in the fans-shaped body and EPG neurons in the ellipsoid body are synchronously recorded as a fly walks in clean air and then tracks the edge of a 10 mm wide jumping plume. After the first 20 entries to the plume, the plume is periodically jumped 3 mm away from the fly during returns (as indicated by blue arrows) to minimize the chance of incidental returns back to the plume’s boundary. **(c)** Epoch of edge tracking corresponding to the highlighted region in (b). Left: Trajectory of the fly as it exits and re-enters the plume. Middle: EPG and FC2 activity (ΔF/F_0_) for that edge-tracking epoch plotted by wedge in the ellipsoid body for EPG neurons (greyscale) and by column in the fan-shaped body for FC2 neurons (bluescale). Right: Estimated EPG (black) and FC2 (blue) phases for that epoch with an indication of when the fly is in the odor (red) and when the plume is jumped (blue arrows). **(d)** Same as for (c) but for the epoch when the same fly was walking upwind prior to encountering the odor plume corresponding to the lower boxed region in (b). **(e)** Distributions of EPG phases (black) and FC2 phases (blue) during upwind walking (anemotaxis). Thin lines represent individual flies and thick lines represent mean across flies. n=6 flies. Distributions are centered around the upwind direction. **(f)** Distributions of EPG and FC2 phases in the 1 sec prior to returns to the jumped plume. Distributions are skewed toward the plume boundary (crosswind direction to the boundary indicated by vertical dashed line). **(g)** Distributions of EPG and FC2 phases in the 0.5 sec prior to exiting of the plume, corresponding to the epochs immediately prior to the returns shown in (f). Direction to the boundary indicated by vertical dashed line. **(h)** Distribution of the difference between EPG and FC2 phases during jumped returns (as in panel (f), orange) and exits (as in panel (g, blue). Crosswind direction towards the plume boundary indicated by the left dashed line and crosswind direction away from plume indicated by right dashed line. Note FC2 phase leads EPG phase in the direction of the odor boundary during returns. n =6 flies. **(i)** Representative mean EPG phase (black arrows) and FC2 phase (blue arrows) plotted as direction vectors onto the averaged exit and return trajectories for two flies that were tracking different sides of the plume. Transparent lines correspond to individual trajectories. Phase is plotted at 12 evenly spaced points on an interpolated time base (6 points inside the plume, 6 points outside the plume). **(j)** Mean progression of EPG phases (black) and FC2 phases (blue) in the 15 sec prior to when flies returned to the plume after a jump. Thin lines represent individual flies, n=6 flies. Bold lines represent mean across flies. n=6 flies. Crosswind direction towards plume boundary indicated by dashed line.

To gain functional evidence supporting this model, we examined the activity of FC2 neurons. This columnar population encodes a fly’s goal as a stable bolus of activity during bouts of menotaxis^32^, in which a fly walks in a consistent direction relative to a visual landmark, keeping its goal and heading aligned. Moreover, FC2 neurons directly synapse onto PFL3 neurons, an output pathway of the central complex that controls steering by comparing an animal’s current heading with its goal^32,43,48,49^. FC2 neurons receive input from multiple tangential and columnar populations^49^, positioning them to encode distinct goal signals in different behavioral states or contexts. However, whether FC2 neuron activity flexibly represents an animal’s changing goals has not been explored.

To assess whether FC2 neurons reflect a fly’s leaving goal, returning goal or alternate between both, we simultaneously recorded the activity of EPG and FC2 neurons as animals walked in clean air and then initiated edge tracking along an odor corridor. The elevated temperatures associated with two-photon imaging can induce dispersal behavior^32,39,47^, promoting straight trajectories with a strong upwind bias, which significantly reduces the crosswind distance animals walk away from the plume’s edge. Therefore, to encourage flies to rely on their memory of the direction toward the plume, we used the jumping plume in which we periodically shifted the plume’s boundary while flies were on an outside trajectory (**Fig. 6b**). This requires flies to walk further on their return paths (**Extended Data** Fig. 6b), preventing them from incidentally re-encountering the plume’s boundary.

Prior to edge tracking, as flies walked upwind in clean air, we observed that the phases of EPG and FC2 neurons remained largely aligned (**Fig. 6d,e**), matching their relationship during menotaxis^32^. During edge tracking, however, the phase of FC2 neurons frequently diverged from the EPG signal **(Fig. 6c**). Notably, the FC2 phase tracked a fly’s current heading inside the odor and as it initially exited the plume, but began to point back in the direction of the plume several seconds before the animal turned towards it on return trajectories (**Fig. 6c,f-j**). The lag between the emergence of this FC2 signal and the subsequent steering of the fly towards the plume (**Fig. 6j**) resembles the time course of the corrective steering flies display when an arbitrary goal angle is experimentally imposed on this columnar population using targeted optogenetics^32^. FC2 neurons thus appear to selectively signal a memory-dependent goal of the entry angle that could guide their returns to the plume’s boundary.

## Discussion

Odors serve as essential navigational cues for many species despite carrying no inherent directional information. Animals must therefore integrate chemical cues with spatial signals to track an odor to its source, a necessity thought to act as a potent force shaping the evolution of navigational circuits^57,58^. Indeed, the close evolutionary relationship of ancestral forebrain circuits, such as the piriform and entorhinal cortex in mammals^59,60,61^, and the extensive connectivity between the mushroom body and central complex in insects^49,55,62^ exemplify how diverse species have evolved the ability to store memories of their odor experience and bind these to internal representations of space.

In this study, we explored how insects rely on memory mechanisms to navigate a chemical landscape. By developing methods to gain precise experimental control over a fly’s olfactory experience, we show that flies leave the plume and return to its edge repeatedly for meters. While this strategy is reminiscent of prior descriptions that insects can track a surface plume by “feeling for its edge”^17^, by experimentally defining and manipulating the location of the plume’s boundary, we revealed that it is in fact stored as a remembered olfactory landmark. Our analyses suggest a model for how the plume’s edge is endowed with salience: each time a fly enters or exits the odor corridor, it dynamically updates and stores a memory of its current heading direction relative to the wind. During edge tracking, flies alternately rely on the integrated memory of their exit and entry angles to guide their trajectories away from the plume and back to its boundary. Consistent with this, we find that FC2 neurons, which have been shown to encode a fly’s allocentric goal^32^, signal the direction of the plume’s boundary during returns.

Flies thus appear to navigate structured chemical landscapes using a vector-based strategy similar to what central place foragers, like ants and bees, use to home to their nests^3,5^. However, unlike a fixed nest location, the boundary of an odor plume is not stationary. Consequently, edge-tracking flies need only store the relative direction to the plume’s boundary rather than its specific position, simplifying memory requirements. Consistent with this, flies do not revisit the site at which they exited the plume but rather bias their trajectories in the direction of the plume’s boundary, as they continue to advance along its length. While the wind serves as an obvious directional cue that flies could use to progress up a vertically oriented plume, we find that flies make similar progress along plumes oriented orthogonal to the wind, by using consistent entry and exit angles that propel them in a chosen crosswind direction. These results underscore that odors themselves are not the ultimate ‘goal’ during plume tracking but rather serve as chemical signposts that an animal can use to track towards the odor’s source. Edge tracking is therefore reminiscent of the route following strategy ants display when navigating through dense visual landscapes^63^, in which they use a sequence of stored local vectors to guide them back to their nest, each associated with a distinct visual snapshot along their route. Serially sampling sensory landmarks, whether olfactory or visual, thus appears to be a shared component of the insect navigational toolkit^13^.

For memories to have predictive value, their timescales must be aligned with the stability of the environment. Once formed, the memory of the plume’s edge appears to be maintained over tens of seconds, as revealed by the persistent directional bias flies display once the plume vanishes. The entry angle memory can nevertheless be updated rapidly over additional odor encounters, such that flies can flexibly adapt to plumes oriented in different directions relative to the wind. While we focused on navigation in a highly-structured chemical landscape to facilitate mapping of a directional memory to plume encounters, such rapid and flexible updating suggests a similar mechanism could be relevant even in the context of more turbulent plumes. The replay of an odor sequence, where a fly encounters boluses of odor uncoupled from its behavior, offers insight into how vector-based navigation could shape a fly’s trajectories even in more chaotic sensory environments^64^. Modeling of edge tracking behavior, however, indicates that memory updating is not instantaneous, but reinforced across successive odor encounters. This suggests the possibility that in a turbulent plume, where odor filaments arrive from erratic directions, the lack of coherence in an animal’s heading direction at odor onset and offset may erode the strength of any angular memory. If the spatial structure of odor encounters is truly stochastic, flies may resort to more reflexive strategies or those that rely on different forms of short-term memory, such as using plume intermittency to guide navigation towards the source^2,64^. Different tracking algorithms are therefore likely to be engaged depending on the coherence and stability of the environment.

During edge tracking, flies encounter a continuous odor corridor yet, behaviorally, create a temporally intermittent experience of the plume by sampling along its edge. One potential reason flies engage in edge tracking is to minimize sensory adaptation^65–67^. However, flies appear relatively indifferent to the concentration of odor they encounter along the longitudinal axis of the plume, suggesting sensory adaptation is unlikely to be the primary driver of this behavioral strategy. Rather, we propose that the plume’s lateral boundary holds special significance, because as the site of the steepest concentration gradient in our virtual environment, it offers maximal information about the structure of a plume^68^ even if it shifts. Consistent with this, artificial recurrent neural networks trained to track a shifting plume adopt a plume-skimming strategy that closely mirrors edge tracking^69^, demonstrating that following the envelope of a plume rather than the wind direction represents a particularly robust navigational strategy. Exploiting environmental contrast by following the steepest sensory gradient has been proposed to underlie navigation in many biological contexts, from the paths of axonal outgrowth^70^ to sonar localization of bats^71^, suggesting edge tracking may represent a general algorithm.

Our data suggest that edge tracking depends on the core navigational circuitry of the central complex, with wind serving as an essential anchoring cue to allow flies to orient themselves in space. While our switching state space model was derived only from behavioral data, it displays striking correspondence to neural components within the central complex that are poised to integrate olfactory and spatial variables and assign meaning to directions in space. In particular, tangential neurons within the fan-shaped body responsive to odors^55^ are positioned to provide modulatory signals onto columnar neurons that encode spatial variables such as an animal’s current heading or traveling direction^46,47,49,50^ to write in the exit or entry angle memory. The alternation between leaving and returning states suggests that two distinct angular memories may be held in parallel and sequentially accessed as a fly weaves in and out of the plume. Consistent with this, a correlate of the entry angle memory emerges in FC2 neurons as animals return to the plume. Notably, we found no evidence that FC2 neurons convey signals related to the exit angle, suggesting additional columnar subsets may signal the goal direction during the leaving state. Alternatively, since the exit angle is predominantly in the upwind direction, an explicit memory-based mechanism within the central complex may not be required. In the leaving state, odor-responsive tangential neurons could suppress the activity of neurons encoding the entry angle memory, to allow flies to reflexively surge upwind^9,10^. Either way, our results underscore how animals rely on dynamically recruited goals to guide their trajectories as they sample along the plume’s boundary.

The alternation between the leaving and returning state could tune the trajectories of an individual^5^, potentially giving rise to the striking variation in the efficiency of edge-tracking trajectories we observe, where some animals rapidly revisit the edge, adhering closely to the plume’s boundary while others take more exploratory circuitous routes (**Extended Data** Fig. 6d). Tangential neurons innervating different layers of the fan-shaped body integrate from distinct ensembles of lateral horn and mushroom body pathways^49,55^, which could allow them to dynamically regulate the strength of memory updating or the timescales of the leaving and returning states depending on the innate or learned valence of an odor or other contextual cues.

The central complex is thought to underlie navigation in many different insect species, from the long-distance migrations of monarch butterflies to ants homing to their nest^3,5,72^. Here we reveal how flies dynamically update angular memories to store direction, not distance, to a plume, exemplifying a basic building block for these more complex navigational feats.

## Methods

### Fly Husbandry

Flies were maintained at 23-25°C and 60-70% relative humidity under a 12-hour light/dark cycle. The quality and composition of the fly food was an important factor for eliciting robust edge-tracking behavior in tethered animals, particularly during imaging experiments. We found that animals raised for several generations on Wurzburg food^50^ more consistently exhibited robust edge tracking than those raised on standard cornmeal-agar-molasses food. All behavioral experiments were conducted using starved flies which were removed from food and placed in vials containing only a water-soaked KimWipe or cotton flug for 16-24 hours before tethering. For optogenetic experiments, flies were reared in complete darkness. Two days prior to an experiment, 1–2-day old flies were transferred to a food vial containing 0.4 mM all-trans-retinal (Sigma #R2500). 16-24 hours before an experiment flies were removed from food and placed in a vial containing a Kimwipe soaked in 1-2 ml of 0.2mM all-trans-retinal in water.

### Detailed fly genotypes

For all behavioral experiments examining edge tracking of an odor plume (Fig. 1, Fig. 2, Fig. 4, Fig. 5, Extended Data. 2, Extended Data. 3, Extended Data. 5, Extended Data. 7, Extended Data. 8, Extended Data. 9), we used:

- Canton-S.

For functional recording and perturbations of EPG neurons (Fig. 3), we used the SS00098 EPG split line:

- 19G02-p65AD ; R22E04-DBD /UAS-GCaMP7s
- 19G02-p65AD; R22E04-DBD /UAS-GtACR1-EYFP

For functional recording of FC2 neurons (Fig. 6) we used:

- VT065306-AD/UAS-syt-jGCaMP7f; VT029306-DBD /60D05-Gal4

For optogenetic activation of olfactory sensory neurons (Extended Data. 4), we used:

- UAS-ChrimsonmVenus/Orco-GAL4; +/+; +/+
- w^1118^ UAS-CsChrimson.mVenus; +/+; +/+ Orco-GAL4, w*; +/+; +/+

*Drosophila* stock sources:

EPG Split Line: 19G02-p65ADZp (in attP40) ; R22E04-ZpGdbd (in attP2) Bloomington *Drosophila* Stock Center (BDSC 93169); FC2 Split Line: VT065306-AD; VT029306-DBD (gift from G. Maimon); R60D05-Gal4 (BDSC 39247); 10XUAS-sytGCaMP7f (attP2) (BDSC 94619); 20XUAS-IVS-CsChrimson.mVenus}attP18 (BDSC 55134); Orco-GAL4.C}142t52.1, w[*] (BDSC 23909); UAS-GtACR1.d.EYFP}attP2 (BDSC 92983); 20XUAS-IVS-jGCaMP7s}VK00005 (BDSC 79032)

### Fly tethering and dissection

All assays were performed using 1–5-day old female flies. Flies were briefly anesthetized (<10 sec) using CO_2_ and tethered to a custom-milled fly-plate similar to what has been previously described^73^). Flies were mounted to the fly-plate using a strand of hair or a single paint brush bristle, which was used to secure their head and subsequently their body to the plate before gluing the eyes and thorax using UV curable glue. In all assays, the filament was removed after successful tethering and flies were placed in a dark, climate-controlled space (25 °C, 40-60% relative humidity) to recover for 15-30 minutes prior to the start of the experiment. Flies were then transferred to the closed loop apparatus and allowed to walk freely on the ball for at least 15 minutes prior to experimentation.

For functional imaging experiments fly preparation varied accordingly: After tethering, the proximal portion of the extended proboscis was glued to minimize movement during recording while allowing the distal portion of the mouthparts to move freely during experiments. Flies were provided a recovery period of 30-120 minutes after tethering. After the recovery period, the fly-plate was then filled with saline (108 mM NaCl, 5 mM KCl, 2 mM CaCl_2_, 8.2 mM MgCl_2_, 4 mM NaHCO_3_, 1 mM NaH_2_PO_4_, 5 mM trehalose, 10 mM sucrose, 5 mM HEPES sodium salt, pH 7.5 with osmolarity adjusted to 275 mOsm). The cuticle covering the posterior portion of the brain was then cut using a 30-gauge needle and removed using forceps to facilitate optical access to central complex structures. Obstructing trachea was removed taking care to not damage the antenna or the antennal nerves. Flies were subsequently transferred and allowed to walk on the ball for at least 15 minutes.

### Olfactory environment preparation

A virtual olfactory environment was created for walking tethered flies leveraging a previously described^16^ closed-loop olfactory system with the addition of custom python scripts that allowed for 2D odor-plume rendering.

#### Tethered locomotion

For tethered locomotion experiments, a spherical treadmill was designed based on previous studies. A 6.0 - 6.5 mm diameter ball was shaped from LAST-A-FOAM FR-4618 (General Plastics) by a custom-made steel concave file. The ball rested in an aluminum base with a concave hemisphere 6.75 mm in diameter with a 1-mm channel drilled through the bottom and connected to an airflow. The ball was recorded at 60–61 fps using a Point Grey Firefly camera (Firefly MV 0.3 MP Mono USB 2.0, Point Grey, FMVU03MTM-CS) with Infinity Lens (94-mm focal length) focused on the ball, with illumination from infrared LED lights. Ball rotation was calculated in real time using FicTrac software^74^ running on computers with ≥3-GHz processor speeds.

#### Closed-loop arena

The heading of the fly, as calculated by FicTrac, was transmitted to a RaspberryPi 4 via serial port. Custom Python code was used to translate heading into tube position controlled by motors described below. The closed-loop air delivery system was custom designed using OnShape (www.onshape.com) and 3D printed using VisiJet Crystal material at XHD resolution in a 3D Systems ProJet 3510 HD Plus. O-ring outside dimension (OD) and inside dimension (ID) gland surfaces were designed with excess material for printing and then manually modified on a lathe for improved RMS (surface) finishing. 360° tube rotation was driven by a bipolar stepper motor (SureStep DC integrated NEMA 17 stepper) controlled through an integrated driver and coupled by a Dust-Free Timing Belt (XL Series, 1/4” width (McMaster-Carr, 1679K121, trade no. 130 × L025) to the rotating tube system, which rotated mounted on an Ultra-Corrosion-Resistant Stainless Steel Ball Bearing (3/4” shaft diameter, 1–5/8” OD, McMaster-Carr, 5908K19). The air channel was kept airtight using oil-resistant O-rings (1/16” fractional width, dash no. 020, McMaster-Carr, 2418T126). Motor rotation was measured by a rotary encoder (CUI Devices, AMT10 Series) that was used to correct for skipped steps.

#### Airflow and Odor Delivery

Odor delivery was achieved by directing a continuous stream of humidified clean air through a 2-mm-diameter tube made of VisiJet Crystal material directed at the fly’s antennae. An anemometer (Kanomax 6006-DE) was used to ensure that airspeeds reaching the fly were maintained between 15 and 25 cm/s. Air to the system was first passed through a charcoal filter and humidified by bubbling the air through a DI water reservoir. The air flow was then split between the spherical treadmill and 3 Mass Flow Controllers (MFC, Alicat, MC-Series, 1000SCCM). Downstream of each MFC, air passed through the headspace of odor vials containing either Apple Cider Vinegar (Heinz), 1-octanol or DI water. The ratio of air entering each vial was dependent on the fly’s position relative to the odor plume such that when a fly is outside the boundaries of the plume, 100% of the air is directed into the water vial; when the fly is within the boundaries of the plume, the air is directed at an experimenter-determined ratio between the water vial and the odor vials. Air streams coming from the odor vials then merge with a Y-connector before entering the air delivery system. Thus, by controlling how much of the air passes through each vial, the MFCs control the total odor concentration of the airstream reaching the fly. The mass flow controllers were controlled using a RaspberryPi 4 running custom python scripts.

#### Optogenetic Stimulation

All optogenetic experiments were conducted on flies with intact cuticles. Flies were reared in the dark, transferred to retinal food, and tethered as described above. For optogenetic experiments, a fiber-coupled LED was controlled by a T-cube LED driver (Thorlabs, LEDD1B) to deliver light to the fly’s head in closed-loop with its behavior. For activation of Orco+ sensory neurons, a 660 nm (red) LED (Thorlabs M660FP1) was turned on (0.863 μW/mm^2^) when the animal was inside the fictive odor plume. For EPG inhibition experiments, a 530 nm (green) LED (Thorlabs M530F2) was turned on (0.752 μW/mm^2^) for the duration of the entire trial.

#### Closed loop parameters

Behavioral measurements (sampled at 60 Hz) were determined by the rotation of the foam ball and obtained from FicTrac (x position, y position, heading, roll, pitch and yaw). These were saved alongside MFC flow values, experimental variables (odor on, LED on) and a timestamp in a single .log file.

#### Edge-tracking behavioral assays

Tethered behavioral assays were conducted in a dark, climate-controlled room (25-27 °C, 40-60% relative humidity). Additionally, the closed-loop olfactory system was enclosed in a black tarpaulin as a further shield from other potential light sources, such as computer monitors and indicator lights on system hardware. To prevent odor build-up in the enclosed space a vacuum line (60 cm/s) was placed at the back of the enclosure as an exhaust. Unless otherwise stated, all experiments started with a 2–5-minute baseline period in which flies walked in clean air delivered in closed loop. At the end of this baseline period, flies were placed at the center of the odor plume’s short axis, with the exception of the 90-degree plume in which flies were placed at the center of the odor plume’s long axis. The geometry of the plume (0, 45-, 90-degree) was determined prior to the start of the experiment. Unless otherwise stated, each plume’s short axis measured 50 mm and long axis 1000 mm. For EPG and FC2 imaging experiments the plume’s short axis was 10 mm. For the jumping plume experiments in Fig. 2h, 3e and Extended Data. Fig 5h, 6f, the edge of the plume was shifted 20 mm in the direction opposite to the fly’s heading as the fly exited the plume. Due to the propensity of flies to engage in straighter trajectories due to the heat of the two-photon laser^39^, during functional imaging of FC2 neurons (Fig. 6), once the fly had made 20 plume entries, the plume was jumped 3 mm on every third subsequent entry. For all experiments, trials were terminated if one of three criteria were met: 1) The fly traveled the length of the long axis of the plume provided (1000 mm); 2) The fly traveled over 500 mm perpendicular to the plume’s edge; 3) The fly traveled 100 mm downwind from its starting position on the plume.

A small fraction of tethered flies (<10%) did not acclimate to walking on the ball or did not show a behavioral response to ACV (by turning upwind and increasing their speed) and were excluded from the study.

#### Odor concentration gradient plumes

Gradient plumes were created with the same dimensions (50 mm by 1000 mm) but featured one of three odor concentration gradients:

#### Vertical plume with increasing upwind odor concentration

Odor concentration increased linearly in the upwind direction, starting at 10% ACV at the plume’s downwind end (0 mm) and reaching 100% ACV at the upwind end (1000 mm). **Vertical plume with decreasing upwind odor concentration:** Odor concentration decreased linearly in the upwind direction, starting at 100% ACV at the downwind end (0 mm) and diminishing to 10% ACV at the upwind end (1000 mm). **90-degree plume with crosswind odor concentration gradient:** Odor concentration varied linearly along the lateral axis, increasing from 10% to 100% ACV in one crosswind direction or decreasing from 100% to 10% ACV in the opposite direction. Flies were initially positioned at the interface between these two gradients, allowing them to track up or down the concentration gradient in the crosswind direction.

#### Optogenetically generated fictive odor plumes

Fictive odor plumes were the same dimensions as a vertical odor plume (50 mm by 1000 mm). Flies were positioned in the center of the plume’s short axis and optogenetic stimulation was provided via a 660 nm LED whenever the fly was within the plume’s boundaries.

#### Disappearing plumes

Flies tracked a constant concentration plume for 10 minutes. After this period, during their first trajectory outside the plume, the plume was removed, leaving the fly walking in closed-loop wind without further olfactory input.

#### Replay experiments in which the odor sequence was replayed back to the fly in open loop

For odor replay assays, we allowed flies to track a 90-degree plume for 10 minutes and recorded the timestamps of odor onset and offset in a separate log file. This file was used to generate the temporal sequence of odor pulses delivered during the replay epoch of the experiment which was initiated 10 sec after the edge-tracking epoch back to the same fly. The same temporal sequence of odor pulses was also presented to a naïve fly that had never edge tracked. Consequently, for every replay experiment, we collected data from one naïve fly and each naïve fly received a distinct temporal sequence of odor pulses defined by the matched replay experiment. During replay, wind was delivered in closed-loop but the animal’s fictive position had no impact on the timing of odor delivery.

#### Operant training paradigm

The operant training paradigm consisted of three phases:

1. Initial edge-tracking phase: Flies were allowed to track a 45-degree plume. After tracking for 250 mm along the plume’s edge, the plume disappeared as flies were on their next outside trajectory.
2. Operant training phase: During training, a fly’s average heading and speed was continuously calculated over a 2 sec sliding window. Flies triggered the delivery of odor when their average heading was in a 45-degree range of entry angles (either -90° to -135° or 90° to 135° depending on whether training was in the same direction as the initial plume or in the opposite direction, see **Extended Data** Fig. 11a) while sustaining an average speed of > 2 mm/sec . Odor was delivered for a minimum of 2 sec and terminated when animals oriented in the 90-degree range of upwind exit angles (-45° to 45°) while sustaining an average speed of >2 mm/sec. The number of training epochs (e.g. one or five odor stimulation periods) during this training phase and whether they were in the same or opposite direction as the initial plume segment was determined prior to the start of the experiment.
3. Test edge-tracking phase: After completing the predetermined number of training epochs, flies were positioned at the center of the short axis of the test plume, oriented opposite to the direction of the 45-degree plume they had tracked during the initial phase. Note that after flies completed this training phase, they had to enter the test odor using the same heading as used for the training phase and so effectively had an additional epoch of reinforcement. In the test phase, the plume extended 20 mm below the fly’s starting position to increase the chances that they successfully entered the test plume.

### Analyses of edge tracking behavior

#### Trajectory averages (Fig. 1e, 2e-h, Extended Data Fig. 5d, 6a)

To construct average trajectories, x and y coordinates for each outside or inside bout were upsampled to contain 10,000 points through linear interpolation between data points using the pandas library in Python. This allowed us to average x and y coordinates regardless of the length of the inside or outside bout. Upsampled x and upsampled y coordinates were averaged to produce one set of (x,y) coordinates for each animal. To construct an average across animals, the averages generated from individual animals were averaged together.

#### Entry and exit angles

Entry and exit angles were determined by calculating the mean traveling direction in the 0.5 sec prior to and 0.5 after crossing the plume’s boundary.

#### Analysis of inbound and outbound segments (Fig. 1f, Extended Data Fig. 5d)

The outside trajectories of flies tracking a vertical plume were simplified into straight line segments using the Ramer-Douglas-Peucker (RDP) algorithm,^75^ which simplifies a set of x, y coordinates by iteratively reducing the number of points in the trace. The parameter ε determines the maximum allowed distance between the simplified and original trajectories and was 1 mm. These trajectories were then averaged using the linear interpolation method described above providing a single average trajectory for each animal. From this average trajectory, “outbound” (away from the edge) and “inbound” (back towards the edge) path lengths were calculated as follows: (1) outbound pathlength was calculated as the pathlength from the point of exiting the plume to the farthest point orthogonal to the plume’s edge and (2) inbound pathlength was calculated as the pathlength from the farthest point orthogonal to plume’s edge back to the plume’s boundary. These three points – plume exit point, farthest point and plume entry point – define a triangle, the interior angles of which define inbound and outbound angles as indicated in Fig. 1f. For analyses of simulated inbound and outbound trajectories in a random search model (Extended Data Fig. 5d), we took the total number of outside bouts for each fly and then randomly selected the same number of outside simulated outside trajectories. From this collection of random trajectories, we used the method above to generate an average trajectory and then calculated outbound/inbound lengths.

#### Anemotaxis definition (Fig. 6e)

Trajectories of animals prior to ACV exposure (pre-air period) were divided into anemotaxis bouts, defined in a similar manner to menotaxis bouts in (Mussels-Pires et al 2024). Animals’ trajectories were simplified into straight line segments using the Ramer-Douglas-Peucker algorithm (epsilon = 10 mm). Segments of length greater than 50 mm were assigned as anemotaxis bouts.

#### Random-walk model (Fig. 1g, Extended Data Fig. 5)

We simplified the outside trajectories of 40 flies tracking a vertical plume using the Ramer-Douglas-Peucker algorithm (epsilon=1 mm) and extracted the straight-line distances (run lengths) that flies traveled between successive turns and the angles between consecutive straight-line vectors (turn angles) to capture the changes in heading direction at each turn and stored each in a separate library.

We simulated random-walk trajectories by randomly sampling from the two empirical distributions of run lengths and turn angles. Simulated flies started at the plume’s edge and the initial run length was assigned a heading that was randomly drawn from the initial segment heading directions from the plume (**Extended Data** Fig. 5b). At each step:

1. A run length was randomly sampled from the empirical distribution of run lengths.
2. A turn angle was randomly sampled from the empirical distribution of turn angles.
3. The simulated fly moved forward by the run length and then adjusted its heading by the turn angle.

The simulation continued until the simulated fly either re-encountered the plume edge or exceeded a maximum distance traveled 500 mm in the direction perpendicular to the plume’s edge which is the same maximum distance flies were allowed to travel in experiments before a trial was terminated.

To reflect the natural tendency of flies to walk upwind even in the absence of odor cues^16^, we modified the random-walk model to include an upwind bias. We adjusted the turn direction based on a sinusoidal probability function whose magnitude scaled between 0 and 1 [value *a*]. When *a*=0, the probability of turning left or right is the same for all heading directions. As the values of *a* increase from *a*=0 to *a*=1, the fly becomes progressively more biased to turn left when it is facing right, and to turn right when facing left. The value of *a* was selected to match that of real flies (**Extended Data** Fig. 5c) mirroring the upwind bias observed in actual flies outside the odor plume.

Given that random exploration outside the plume should not depend on the geometry of the plume tracked, the same model was used to simulate outside trajectories for flies tracking the 45-degree and 90-degree plume, drawing from same empirically-derived libraries of run lengths and turn angles (**Extended Data** Fig. 5). The only difference between the models was the initial heading which was sampled from the initial segment heading directions for the 45- and 90-degree trajectories.

#### Functional imaging

All functional imaging experiments were performed on either an Ultima or Investigator two-photon laser scanning microscope (Bruker) equipped with galvanometers driving a Chameleon Ultra II Ti:Sapphire laser. Emitted fluorescence was detected with GaAsP photodiode (Hamamatsu) detectors. Images were acquired with an Olympus 40X, 0.8 NA or 20X, 1.0 NA objective at 512 pixels × 512 pixels resolution. The laser was tuned to 920 nm for all experiments. The out-of-objective power for all experiments was ∼15mW and never exceeded 25 mW. For volumetric imaging, 3-5 slices were recorded using a piezoelectric Z-focus (Bruker) at a rate of 3-10 Hz.

#### Data alignment

To align the imaging and behavioral data, we used a 3.3V digital pulse from a RaspberryPi to initiate recording of an imaging stack through the PrairieView (Bruker) I/O interface. The RaspberryPi continued to send 3.3V pulses every 10s which were recorded with PrairieView for confirmatory alignment of the imaging and behavioral data. Behavioral variables were captured at 60–61 frames per second, whereas neural activity (fluorescence) was captured at 3-10 frames per second. To align these two distinct time series, a standardized and regular time series was generated for each trace containing 100-ms time bins (0, 100, 200, 300, etc.). Behavioral data was binned at the center of each time point and averaged. Imaging data was upsampled using linear interpolation at the same 100 ms time bins. Custom Python scripts were used to synchronize and log incoming FicTrac variables and outgoing MFC and motor commands, allowing us to align the wind direction, odor concentration, behavioral data and imaging frames.

#### Image registration and ROIs

All images were stored as individual .tiff files which were registered using the StackReg function of the pystackreg library. ROIs were drawn using custom Python scripts, Fij, or a custom MatLab GUI. To quantify EPG neuron activity recorded from the protocerebral bridge, we followed the methods described in (Lyu et al., 2022)^46^ and manually defined each of the 16 glomeruli from the registered stacks corresponding to each imaging plane.

FC2 neuron activity during edge tracking was recorded in the fan-shaped body. EPG neuron activity in the ellipsoid body was recorded synchronously to allow for direct comparison of their phase relationships during an experiment. To quantify FC2 neuron activity, we manually defined the lateral borders of the fan-shaped body, extended these lines down to a point and divided the arc defined by these intersecting lines into 16 wedges. Pixels were assigned to each wedge if they resided within the wedge and within the footprint of the FC2 neurons, defined by masks that were manually drawn in each imaging layer based on their arborization pattern. To quantify EPG neuron activity recorded in the ellipsoid body, we radially divided the ellipsoid body into 16 sub wedges from a manually defined center. Pixels were assigned to each wedge if they resided within the wedge and within the footprint of the EPG neurons, defined by masks that were manually drawn in each imaging layer based on their arborization pattern.

ΔF/F_0_ values were determined by the formula (F-F_0_)/F_0_, where F is the mean pixel value within a wedge/glomerulus and F_0_ is the mean of the lowest 5% of F values.

#### Neuronal phase analysis

Phase was determined by two methods. For recordings of EPG neurons in the protocerebral bridge, phase was calculated as in Green et al. 2017^44^. For each timepoint the Fourier transform was taken on the ΔF/F_0_ activity vector of the 16 glomeruli. Phase was determined as the phase of the Fourier spectrum at a period of 8, the peak periodicity of the power spectrum^44^. For recordings of EPG neurons in the ellipsoid body and FC2 neurons in the fan-shaped body, phase was defined as the population vector average of wedge signals as per Seelig and Jayaraman 2015; Lyu et al. 2022^37,46^. In summary, the 16 wedges of the ellipsoid body and fan-shaped body were assigned angles from -180° to 180°, equally dividing 360° of azimuthal space. Angles were assigned so that ellipsoid body and fan-shaped body wedges were in correspondence as per the connectome^49^. In the ellipsoid body, wedges were counted counterclockwise starting from the most ventral wedge to the right of the midline. In the fan-shaped body, wedges were counted from right to left. For each timepoint, each wedge was assigned a vector with length equal to the ΔF/F_0_ value and direction given by its assigned angle. Phase was the angle of the population vector average of these vectors at each timepoint.

#### Phase nulling

Phase nulled EPG signals recorded in the protocerebral bridge were calculated as previously described^44,46^. At each frame, we rotated the signal by the estimated phase of the activity bump in that frame, such that all bumps were aligned. To compare bumps in and out of odor, we separated frames corresponding to when the fly was in or out of the odor and averaged 122 together with the signal from all these frames to get the average bump amplitude.

#### Phase calibration to allocentric coordinates

The anatomical location of activity bumps in the central complex correspond to azimuthal orientations in allocentric space. However, the mapping from anatomical to external coordinates varies between animals and can remap during an experiment^37^. To produce EPG and FC2 signals aligned to external coordinates, we rotated phase and wedges by an offset at each timepoint such that phase angle 0° was aligned upwind and the central two wedges corresponded to 22.5° to the left and right of the upwind direction (**Fig. 6c**). The offset was the rolling mean (over 2 s) of the difference between the EPG phase and the animal’s heading, since the EPG signal provided a faithful neural correlate of heading relative to the wind (**Fig. 3b**). To rotate wedges, this offset was converted to an integer value varying between -8 and 8 by rounding to the nearest 22.5°. The same offset was applied to EPG and FC2 signals.

### Model

#### Time Discretization

We bin the data into time intervals of 0.2 s and run the behavioral model in discrete times steps with this same interval. Because we are modeling the motion of the fly, we excluded data points when the fly’s speed is less than 1 mm/s.

#### Edge Crossing Events for Memory Update

To avoid spurious events and to account for the finite width of the odor boundary, we define an edge crossing when a fly is on one side of the odor boundary for three consecutive times points and then on the other side for three consecutive time points. We define the fly’s velocity unit vector at the time of the crossing, 𝒗̂, as its velocity averaged over this five-time-step interval, normalized to unit length.

#### State

The model has two states: returning and leaving. When the fly leaves the odor plume in the leaving state, a time interval is drawn from a negative binomial distribution NB(r_leave_, 1−p_leave_) and, after that number of time steps, the state switches to the returning state if the fly is still out of the plume. When the fly enters the odor plume in the returning state, a time interval is drawn from a negative binomial distribution NB(r_return_, 1-p_return_) and, after that number of time steps, the state switches to the leaving state if the fly is still in the plume. Otherwise, no transitions occur. The parameters r_leave_, p_leave_, r_return_ and p_return_ are determined by fitting to data by the procedure describe below.

#### Memory

The model stores 2-d vector memories of plume exit and entry directions. The entry memory vector 𝒎_entry_is updated at the time of each entry by

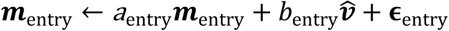

where 𝒗̂ is the entry velocity unit vector defined above, and the two components of 𝛜_entry_ are random variables drawn from a normal distribution with zero mean and variance σ_entry_^2^. The exit memory vector 𝒎_exit_ is updated at the time of each exit by

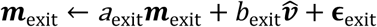

with *v̂* the exit velocity unit vector and 𝛜_exit_ drawn from a normal distribution with zero mean and variance σ_exit_^2^. In simulations, we ignore updates for which the exit 𝒗̂ is more than 60^◦^ away from the upwind direction. This follows the intuition that flies have an innate tendency to exit upwind. Other than during these plume boundary crossings, 𝒎_entry_and 𝒎_exit_remain unchanged. The values of the memory vectors at the beginning of a trajectory depend on unknown previous history. We model 𝒎_entry_(0) as the null vector and 𝒎_exit_(0) with a trajectory-specific Gaussian prior 𝑁(𝝁_0_, 𝜎_0_^2^𝐼) during fitting.

The parameters *a*_entry_, 𝑏_entry_, σ_entry_^2^, *a*_exit_, 𝑏_exit_, σ_exit_^2^, 𝝁_0_ and 𝜎_0_^2^ are determined by the fitting procedure described below. During fitting, nonzero values of 𝜎_0_^2^, σ_entry_^2^ and σ_entry_^2^ allow unexplained variation of latent memories. However, during simulation, we set 𝜎_0_^2^ = σ_entry_^2^ = σ_entry_^2^ = 0 to test the effectiveness of our hypotheses without introducing extra noise into the memories. When simulating from models fitted to a single trajectory, as in Extended Data Fig. 6d, we set the initial exit memory to 𝝁_0_. When simulating from the average fly model (parameters computed from multiple trajectories; **Fig. 5b**), we set the upwind component of the initial exit memory to a fixed value 𝑤_‖_ equal to the median of the upwind components of 𝝁_0_from all training trajectories, and we sample the crosswind component from a uniform distribution over the range from −2𝑤_⊥_ to +2𝑤_⊥_, where 𝑤_⊥_ is the median of the absolute value of the crosswind components of 𝝁_0_from all training trajectories.

#### Velocity update

In the leaving state, velocity is updated at every time step with a decay term, a bias towards the current exit memory, and a noise term,

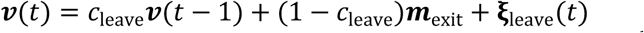

with 𝛏_leave_(𝑡) drawn from a Gaussian distribution with zero mean and variance σ_leave_^+^. Similarly, in the returning state, the velocity update is

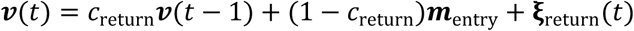

with 𝛏_leave_(𝑡) drawn from a Gaussian distribution with zero mean and variance σ_return_^+^.

#### Variational Inference and Parameter Learning

For each trajectory, we compute the variational distributions of latent variables and trajectory-specific parameters using coordinate ascent of the evidence lower bound^51,52,76^. For r>1, state transitions can be implemented in a Markov chain with an expanded state space, where each original state is augmented with 𝑟 − 1 additional super states. The parameters 𝑟 and 𝑝, for both the leaving and returning states, can then be updated by fitting a negative binomial distribution to samples from the variational distribution of states^76^. For simplicity, we set 𝑟 = 1 during all but the final iteration of the fitting algorithm.

For the average fly model, we fit a single set of optimal parameters (except for 𝝁_0_ and 𝜎_0_^2^, which are still trajectory specific) to latent variable distributions calculated using the above procedure from the 28 trajectories with no less than 30 returns (**Extended Data** Fig. 7c,h). We chose a relatively large threshold on the number of returns to provide a high number of data points for determining the memory update parameters.

For the average fly model, *w*_‖_ = 6.299 mm/s, and *w*_⊥_ = 1.695 mm/s, and, as shown in **Extended Data** Figure 7c:

|  | entry memory | exit memory |
| --- | --- | --- |
| **a** | 0.727 | 0.956 |
| **b** | 0.437mm/s | 0.482mm/s |
|  | return after entry | leave after exit |
| **r** | 2 | 1 |
| **p** | 0.505 | 0.81 |
|  | returning state | leaving state |
| **c** | 0.733 | 0.6 |
| **$\sigma$** | 3.23 mm/s | 3.255 mm/s |

#### Comparison with data

We remove odor changes lasting only 1 or 2 time steps for simulated trajectories before comparing to fly data, which matches the data preprocessed to remove odor changes lasting less than 0.5s.

## Acknowledgements

We thank members of the Ruta laboratory for helpful discussions, R. Mohanta, B. Noro, and S. R. Datta, for comments on the manuscript; P. Stock for initial contributions to the closed-loop olfactory system and providing technical support, P. Mussells-Pires and G. Maimon for sharing the FC2 driver and technical advice. Stocks obtained from the Bloomington *Drosophila* Stock Center (NIH P40OD018537) were used for this study. This study was supported by a Simons Foundation Collaboration for the Global Brain award to L.F.A and V.R., an NIH NINDS grant (R35NS111611) and a Brain Initiative grant (R01NS113103) to V.R., and a Kavli Neural Systems Fellowship to A.F.S. L.F.A. and S.M. are supported by the Simons, Gatsby and Kavli Foundations. N.E. and A.F.S. were supported by a Medical Scientist Training Program grant from NIH NIGMS (T32GM152349) to the Weill Cornell/Rockefeller/Sloan Kettering Tri-Institutional MD-PhD Program. V.R. is a Howard Hughes Medical Institute Investigator.

## Author contributions

A.S. and C.M. designed and performed all behavioral experiments with odor plumes. N.E. performed behavioral experiments using optogenetic activation of sensory neurons. A.S. performed functional recordings of EPG neurons. C.D. performed functional recordings of FC2 neurons. S.M. and L.F.A developed, implemented, and analyzed the model. A.S., S.M., C.D., N.E., J.R. L.F.A and V.R contributed to analyzing and interpreting the data. V.R. wrote the manuscript with input from all the authors.

## Supplementary Information

**Supplementary Video 1:** Representative video of a fly navigating within the closed loop olfactory paradigm. The fly walks on an air-supported ball with a tube carrying clean air/odor yoked to its angular heading.

**Supplementary Video 2:** Video of a representative fly tracking the edge of a vertical plume. Right: view from the back of the fly walking on an air-supported ball with the air/odor tube in front. Middle: Expanded view of fly trajectory as it crosses the border of the fictive odor plume. Right: Trajectory of fly mapped onto 1 meter plume.

**Supplementary Video 3:** Video of a representative fly tracking a jumping plume. Boundaries of plume shown in white lines, which jump 20 mm in the opposite direction of the fly’s heading each time the fly exits. Fly trajectory shown in magenta.

---

**Extended Data Figure 1:**
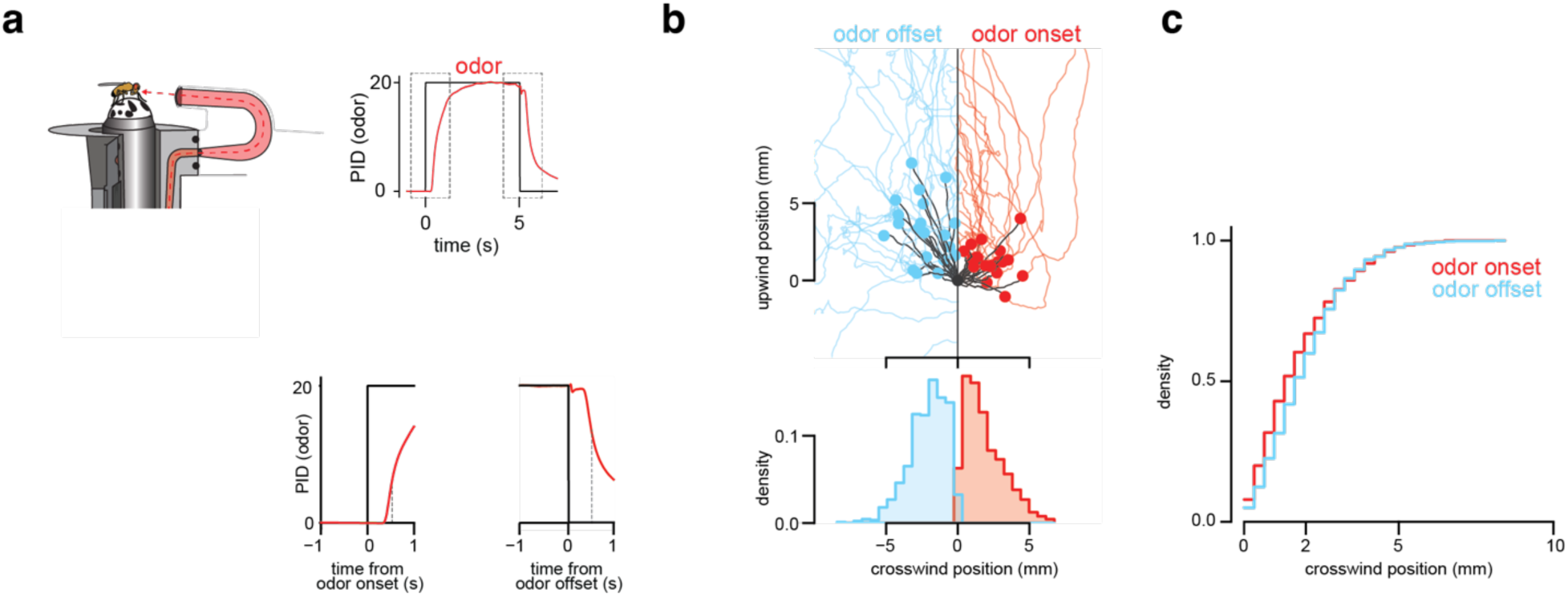
Characteristics of odor stimulation at the plume’s boundary. **(a)** Left: Schematic of closed-loop olfactory system highlighting nozzle yoked to the fly’s heading that provides a continuous stream of clean air, infused with odor (apple cider vinegar) when fly is inside the boundary of the fictive plume. Right: odor concentration profiles measured by PID (photoionization detector) over time at the position of the fly. Top panel shows the full duration of an odor pulse with an onset and decay of approximately 500 msec. Bottom panels show zoomed-in views of the PID trace over the first second from odor pulse onset (left) and offset (right). **(b)** Positional data of a representative fly upon odor onset and offset. Top: the position of the fly for all entries into the plume (red) and exits from the plume (blue). The solid points represent the animal’s position 500 msec after odor onset and offset. Bottom: histogram shows the distribution of crosswind positions of the fly upon exiting the plume (blue) and upon entering the plume (red). Due to the finite time required for the odor to reach the fly as it walks in virtual space shown in (a), the plume’s edge is not perfectly sharp, with a spatial gradient that depends on how fast the fly is walking. **(c)** Cumulative density functions (CDFs) of crosswind positions for flies in response to plume entry (red) and exit (blue). n=40 flies, 731 inside trajectories and 796 outside trajectories.

**Extended Data Figure 2:**
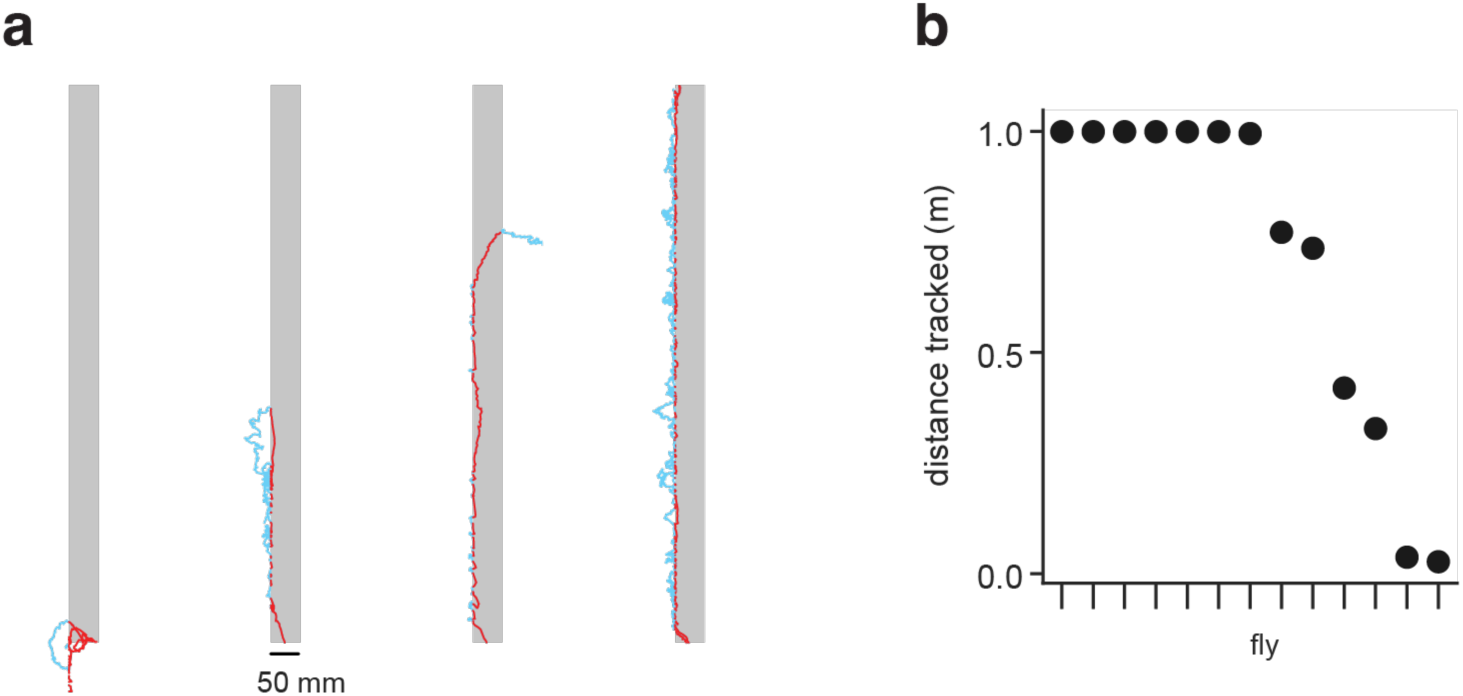
Flies readily edge track along the edge of a vertical plume. **(a)** Representative trajectories of four naïve flies that had never previously encountered an odor corridor tracking a 1-meter vertical plume. Epochs outside the plume are depicted in blue and epochs inside the plume are depicted in red. **(b)** Plume distance tracked by 13 naïve flies. Note that for these experiments, the trial was terminated once a fly tracked more than 1 meter upwind, more than 500 mm in the crosswind direction or more than 100 mm downwind after encountering the plume.

**Extended Data Figure 3:**
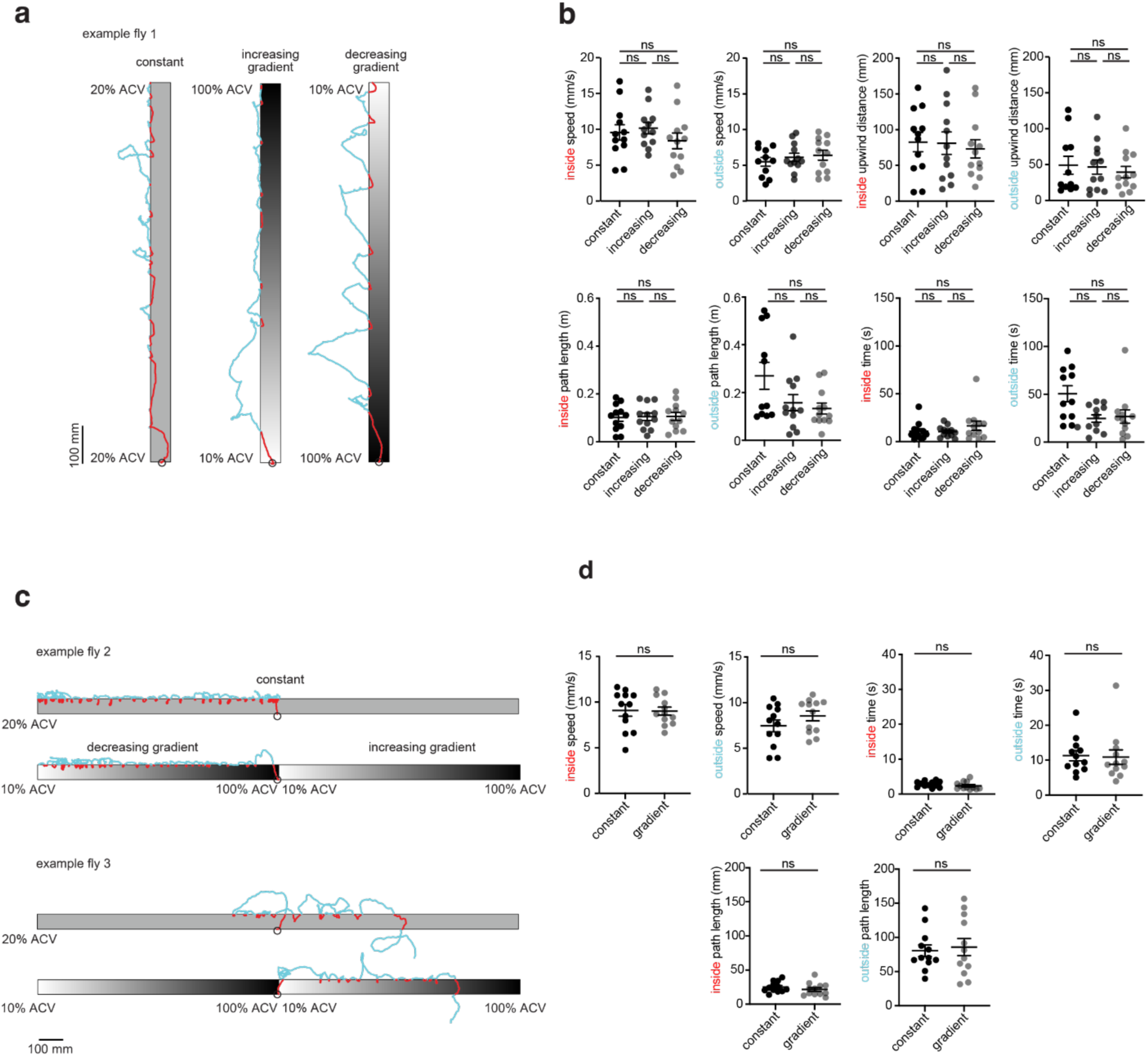
Edge tracking does not depend on the concentration gradient along the longitudinal axis of the plume. **(a)** Representative trajectories of an individual fly tracking an odor corridor of a constant concentration of 20% apple cider vinegar (ACV) (left), a plume with an increasing ACV concentration gradient in the upwind direction (10-100% ACV over 1-meter, middle) and a plume with a decreasing ACV concentration gradient in the upwind direction (100-10% ACV over 1-meter, right). Start of each trajectory is marked with a circle. **(b)** Comparison of average behavioral metrics for the three plumes shown in (a). Each fly navigated a constant concentration, increasing gradient and decreasing gradient plume in a randomized order across flies. Each point represents the average metrics for a single fly, with interindividual average and SEM shown (n = 12 flies). Wilcoxon signed rank test. **(c)** Representative trajectories for two flies that sequentially navigated a constant concentration 90-degree plume (top) and 90-degree plume, in which the fly was positioned at the center and could select to walk up an increasing (10-100% ACV over 1-meter) or decreasing gradient (100-10% ACV over 1-meter, bottom). **(d)** Comparison of average behavioral metrics for the constant or gradient plumes shown in (c). Each point represents the average metrics for a single fly, with interindividual average and SEM shown (n = 12 flies). n.s. not significant, p > 0.05. Details of statistical analyses and sample sizes are given in Table S1.

**Extended Data Figure 4:**
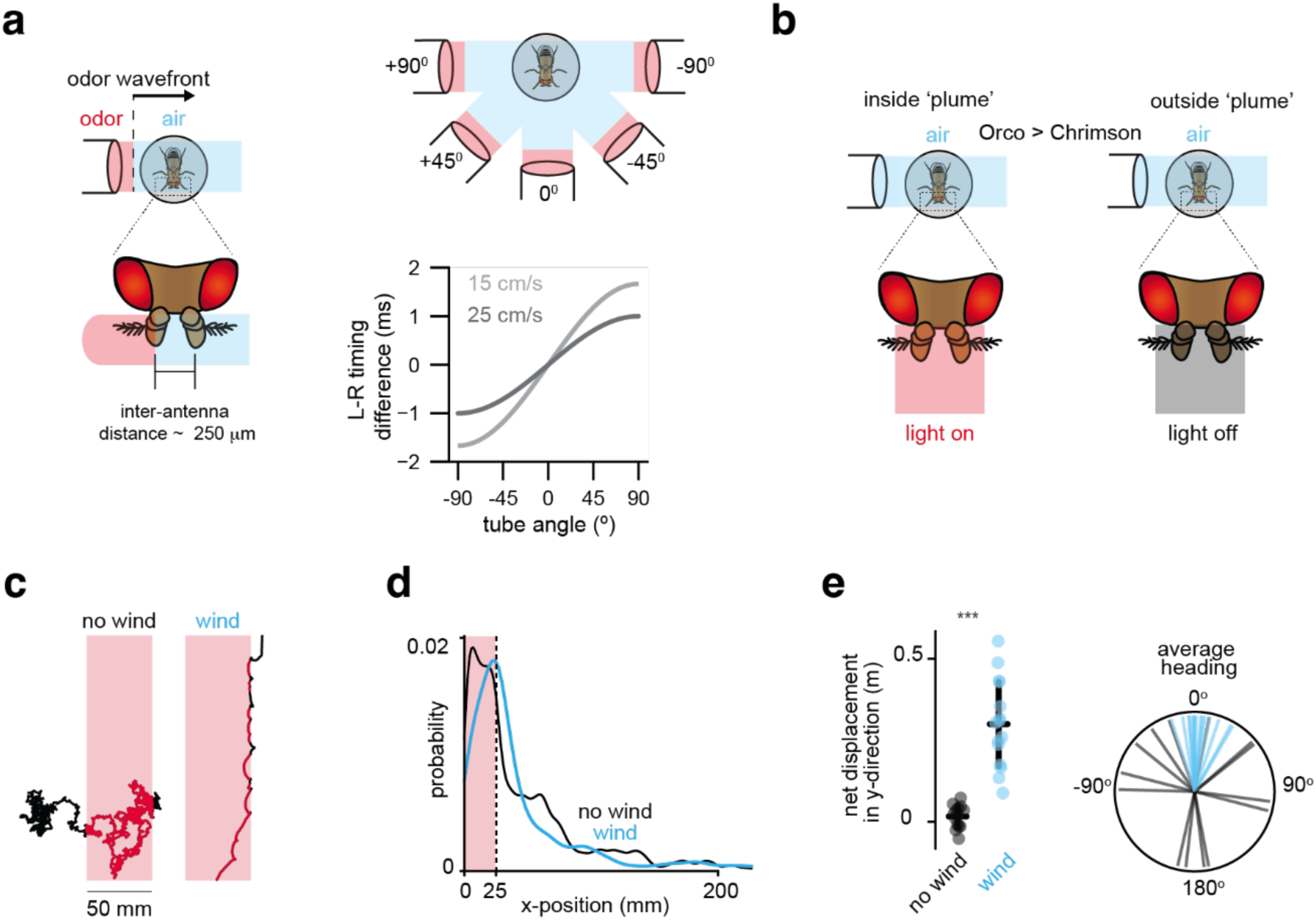
Flies do not appear to require bilateral signals to localize the plume’s edge or track along its boundary. **(a)** Schematic of the experimental setup in closed-loop olfactory stimulation paradigm illustrating the contact timing of an odor wavefront arriving from different angles (+90°, +45°, 0°, -45°, -90°) to each antenna. Given the small spatial distance between the two antennae and the speed of the air flow, the difference in the timing of odor arrival is < 1 ms even when animals are walking in the crosswind direction. **(b)** Schematic of experimental setup for flies navigating a fictive odor plume generated by optogenetic activation of Orco+ sensory neurons expressing CsChrimson. When a fly is inside the boundaries of the fictive plume (left), both antennae are simultaneously stimulated using 660 nm light. Outside the fictive plume (right), the light is off (grey) and there is no stimulation. Wind is maintained in closed loop throughout. **(c)** Representative trajectories of a fly presented with a 660 nm light corridor in the absence (left) or presence (right) of wind. Paths inside the corridor are shown in red while paths outside the corridor are shown in black. **(d)** Probability distributions of the absolute value of flies’ x-positions inside a light corridor (depicted in red) in the presence (blue) or absence of wind (black). n=14 flies (no wind), 15 flies (wind). **(e)** Left: Average displacement in the y-direction for flies presented with a light corridor in the presence (blue) or absence of wind. Right: Average heading inside the light corridor in the presence (blue) or absence of wind (black). n=14 flies (no wind), 15 flies (wind). ***p < 0.001 Details of statistical analyses and sample sizes are given in Table S1.

**Extended Data Figure 5:**
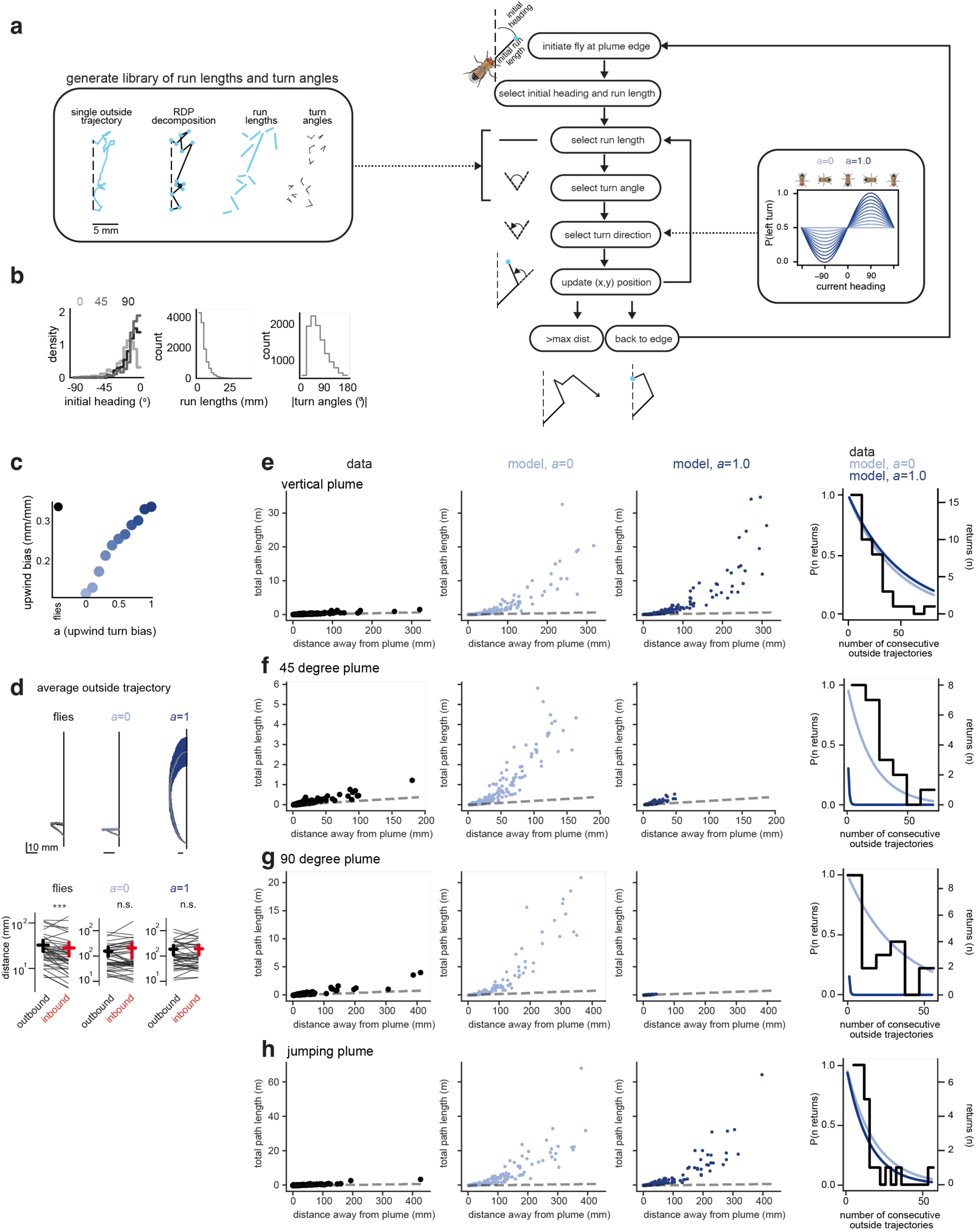
Edge tracking is not the result of random exploration outside the plume. **(a)** Schematic of random-walk model. Left: individual outside trajectories from 40 flies tracking a vertical plume were decomposed into a series run lengths and turn angles using the Ramer-Dougles-Peucker algorithm to generate a library for generating synthetic trajectories with the same underlying statistics. Right: flowchart of how trajectories were simulated from this library. The fly was initiated at the plume’s edge with an initial heading and run length matching the distribution observed during actual fly exits from the plume. At each step, a run length, turn angle, and turn direction was chosen to update the new position (x,y) of the fly. The procedure was repeated until the simulated trajectories returned to the edge or traveled greater > 500 mm orthogonal to the plume’s edge. The turn direction was biased by the wind direction as shown in the inset graph which compares the heading distributions for different upwind biases (*a*=0 to *a*=1.0). **(b)** Left: Distribution plots of initial headings for the three different plume geometries: 0-degree (light grey); 45-degree (dark grey) and 90-degree (black). Middle: distribution run lengths derived from the library of trajectories. Right: distribution of turn angles derived from the library of trajectories. **(c)** Relationship between upwind bias and upwind turn bias (*a*) for simulated and real flies. Upwind bias is defined as the distance traveled in the upwind direction divided by total path length. Note that real flies show an upwind bias of 0.35, which corresponds to an upwind turn bias (*a*=1.0). **(d)** Top: Average outside trajectories for flies (left) and simulated trajectories (right) tracking a vertical plume with different upwind biases (*a*=0, *a*=1.0). Note that in contrast to real flies, simulations lacking an upwind bias (*a*=0) never progress along the length of the plume. Simulations with *a*=1.0 make progress but lack directed returns. Because only simulated trajectories that successfully returned to the plume are included in these averages, they progressed further upwind than real flies but were less efficient as shown in (e). Bottom: Comparison of the inbound and outbound segments of outside trajectories (see Methods), defined as leading away from (outbound, black) or returning to (inbound, red) the point farthest from the plume in the crosswind direction for real flies (replotted from Fig. 1f), and simulated trajectories lacking an upwind bias (*a*=0) and with an upwind bias (*a*=1.0). **(e-h)** Comparison of real and simulated trajectories for different plume configurations: vertical plume (e, replotted from Fig. 1g), 45-degree plume (f), 90-degree plume (g), and jumping plume (h). Comparison of total path length as a function of distance in the perpendicular direction away from the plume for real flies (left column), model with *a*=0 (middle-left column), and model with *a*=1.0 (middle-right column). Right column compares the probability of returning for *n* consecutive outside bouts for real flies in black and random models shown in dark and light blue.

**Extended Data Figure 6:**
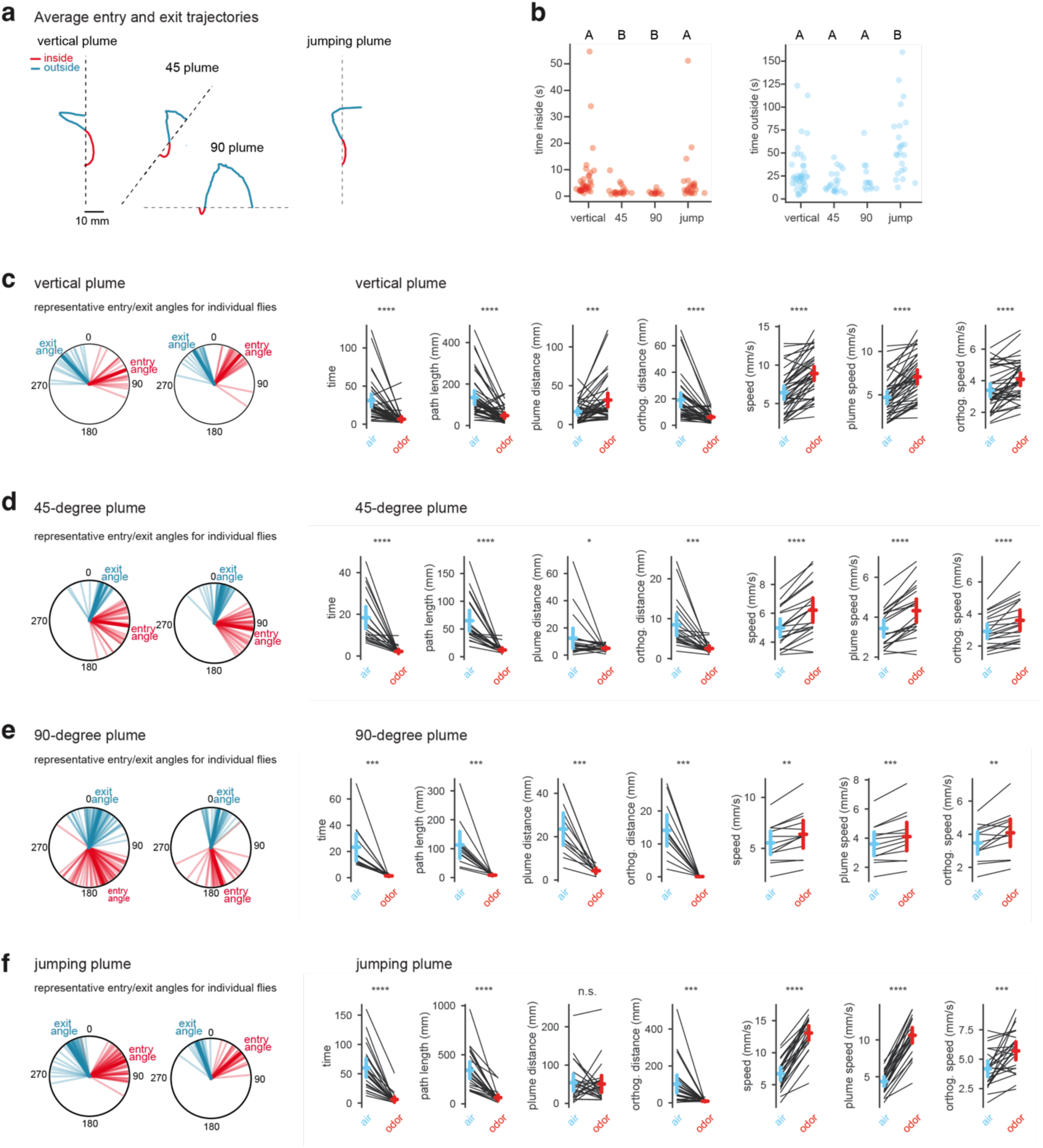
Comparison of edge tracking of plumes with different orientations relative to the wind. **(a)** Comparison of aligned average trajectories for flies tracking indicated plume orientations, n=20-40 flies, replotted from Fig. 2e-h to highlight the variation in the inside (red) and outside (blue) trajectories of flies depending on the geometry of the plume they are tracking. **(b)** Comparison of time spent inside or outside the plume across different plume geometries, with each dot indicating the average of all trajectories for a single fly. n=20-40 flies. Letters denote statistically different groups (p < 0.05). **(c-f)** Left, entry (red) and exit (blue) angles for two representative individuals tracking the indicated plumes shows that flies use a restricted set of angles as they edge track. Right, comparison of behavioral metrics in clean air outside the plume (blue) and in odor (red) for indicated plumes. **(c)** 0-degree (with data replotted from Figure 1); **(d)** 45-degree; **(e)** 90-degree; **(f)** jumping plume. Each data point indicates the average for all inside or outside bouts for an individual trajectory. n.s., not significant, *p < 0.05, **p < 0.005, ***p < 0.001, ****p < 0.0001. Details of statistical analyses and sample sizes are given in Table S1.

**Extended Data Figure 7:**
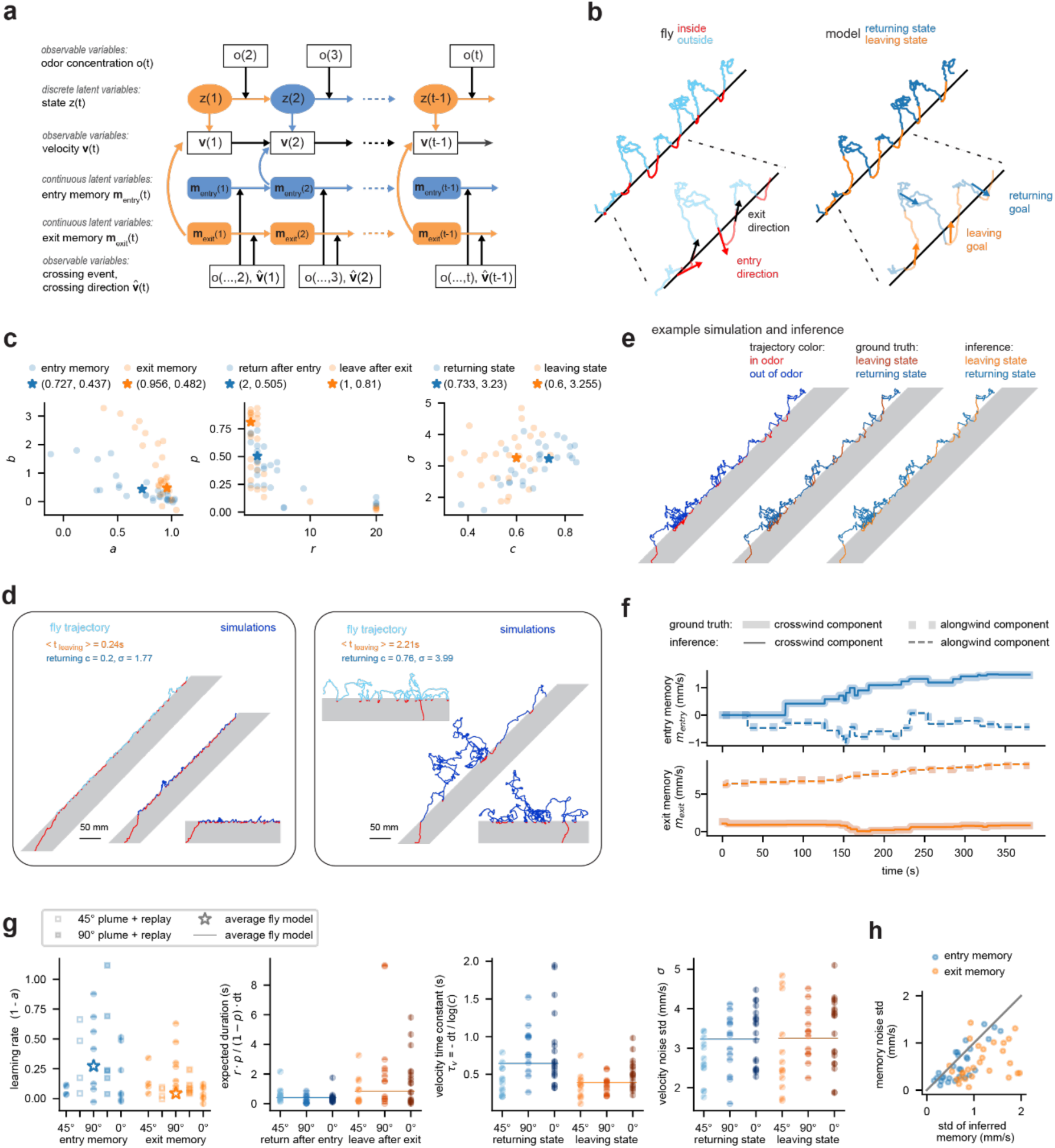
A switching state space model for edge tracking. **(a)** Diagram of the switching state space model used to infer fly navigation states (discrete latent variables) and memories (continuous latent variables). The model also infers transition rates between states and parameters of the memory and velocity update rules. See Methods for details. **(b)** Representative example of a real fly trajectory with navigation states and memories inferred. Left: trajectory is colored according to when the fly is inside the plume (red) or outside the plume (light blue), with arrows indicating entry direction (red) and exit direction (black) as flies cross the plume’s boundary. Right: The same fly trajectory colored according to leaving state (orange) and returning state (blue) with arrows indicating inferred leaving goal (orange) and returning goal (blue). Note that leaving and returning states are not equivalent to times when the fly is inside or outside the odor plume. **(c)** Scatter plots showing the distribution of parameters for the 28 trajectories (dots) from flies that made at least 30 returns which were used to train the model, and parameters for the average fly model (stars), see Methods for details. **(d)** Individual flies display heterogeneity in how closely they track the edge of the plume, reflected in distinct model parameters corresponding to different state transitions and velocity updates. Each box shows the representative trajectory for an individual fly (left, cyan) used to fit model parameters and two simulations (right, blue) for 45- and 90-degree plumes. All bouts inside the odor are shown in red. **(e)** Example inference of navigational states (right) compared with ground truth (middle) for a simulated trajectory. **(f)** Example inference of memories (thin lines) compared with ground truth (thick lines) for the simulated trajectory in (e). **(g)** Parameter distributions across different experiments. The first panel shows the learning rate of 28 training trajectories (trajectories including replay experiment are shown by square markers). The second to last panels show state and velocity parameters for 46 edge tracking trajectories with at least 20 returns. **(h)** Joint distribution of the memory noise parameter and standard deviation of inferred memories for 28 training trajectories. Inferred memories here are the mean of variational distributions fitted to individual trajectories. Standard deviation of inferred memories is calculated as the geometric mean of crosswind component standard deviation and alongwind component standard deviation.

**Extended Data Figure 8:**
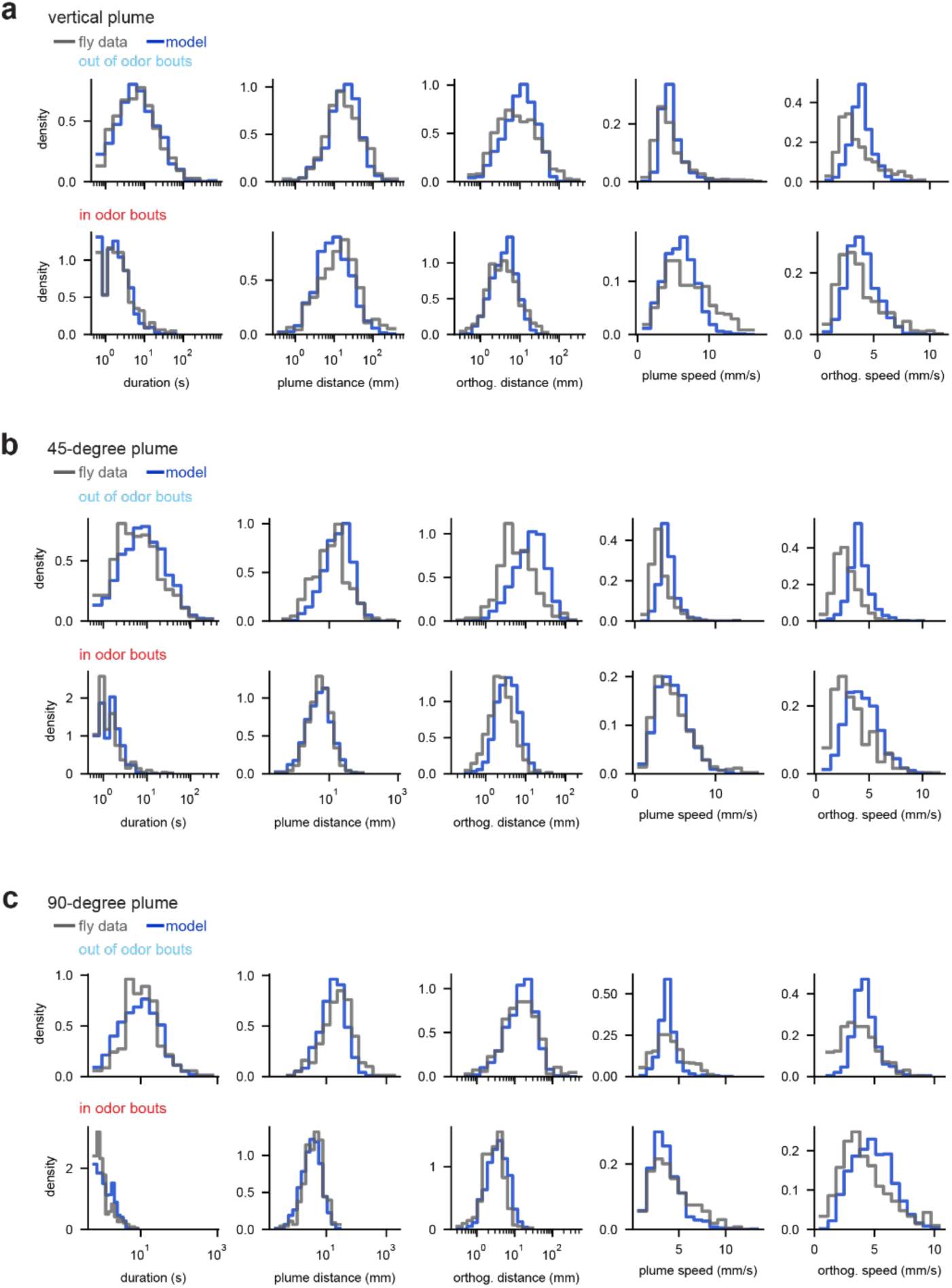
Model captures the average statistics of edge tracking across different plume geometries. **(a-c)** Histograms comparing metrics of real and simulated fly trajectories tracking indicated plume orientations: **(a)** vertical (0-degree) (28 flies, 25 simulations), **(b)** 45-degree (24 flies, 25 simulations), and **(c)** 90-degree (20 flies, 25 simulations). These represent the same set of trajectories as in Figure 5c, d but now divided by plume geometry.

**Extended Data Figure 9:**
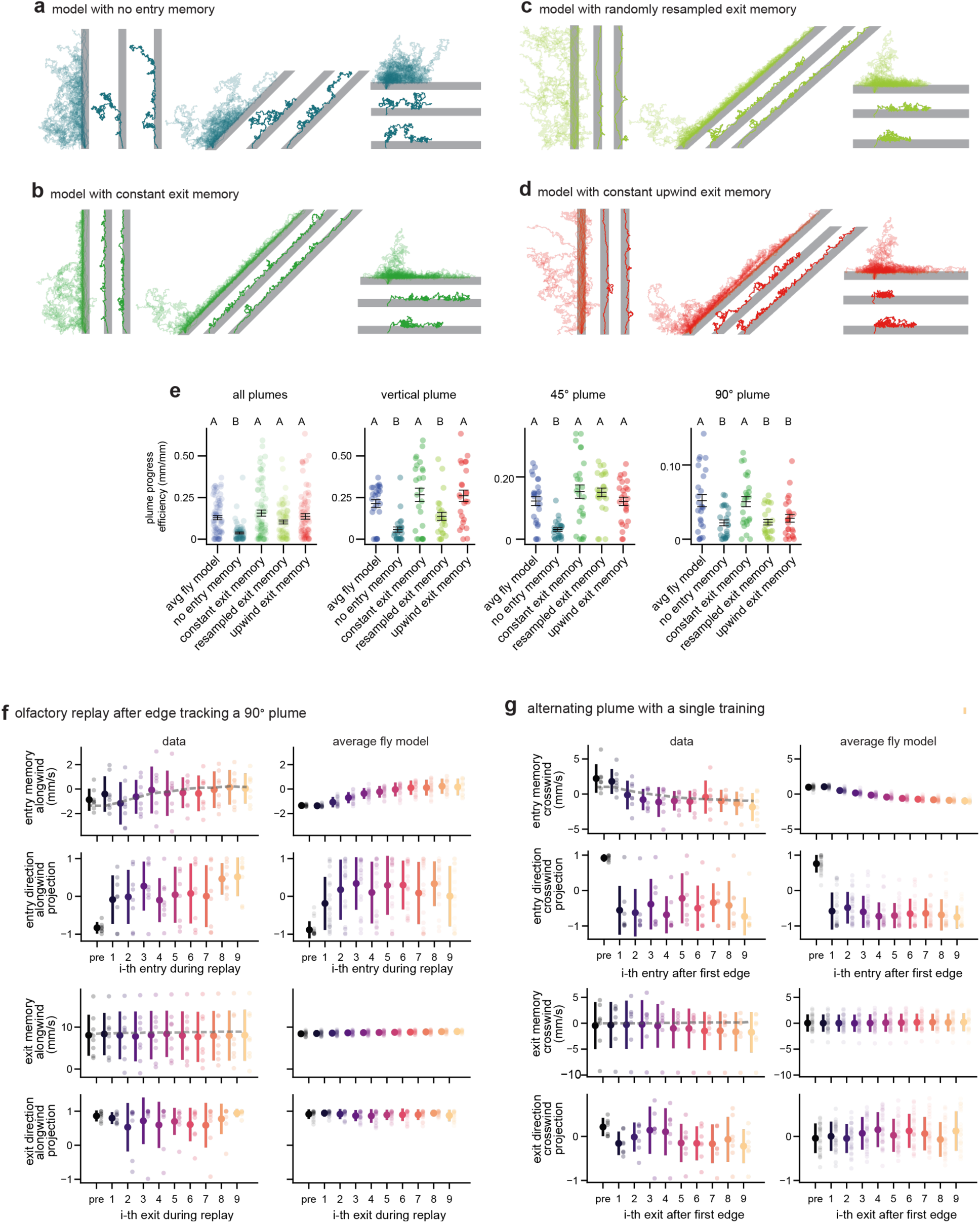
The role of the entry and exit angle memory in the model. **(a)** Simulated trajectories tracking vertical, 45-degree and 90-degree plumes using a model where entry memory is maintained as a null vector. For each plume, two individual representative trajectories are shown (thick lines) along with an overlay of 25 trajectories. **(b)** Same as (a) but for simulated trajectories using a model that has a constant exit memory in the upwind direction with a random crosswind component. **(c)** Same as (a) but for simulated trajectories using a model where the exit memory is resampled from the initialization distribution at each exit. **(d)** Same as (a) but for simulated trajectories using a model with a constant upwind exit memory. **(e)** Plume progress (distance between first and last exit divided by total path length) for indicated model variants. Letters denote statistically different groups (p < 0.05). Details of statistical analyses and sample sizes are given in Table S1. **(f)** Changes of the alongwind component of the entry angle memory, entry direction, exit memory and exit direction during olfactory replay after tracking a 90° plume (see Fig 4b**,c**). Results shown for trajectories with more than 9 entries during replay. Left: Entry and exit direction and inferred entry and exit angle memory for 8 real flies. Inferred memories are the mean of variational distributions fitted to individual trajectories. Dashed gray line indicates average from corresponding model trajectories plotted for comparison. Error bars show standard deviation. Right: Same as at left but plotted for 22 simulated trajectories. For simulations, entry and exit angle memory represent ground truth memory. **(g)** Changes of the crosswind components of entry memory, entry direction, exit memory and exit direction for alternating plume experiment (see Fig 5e**-f**) with a single operant training period. Results shown for trajectories with more than 9 entries after tracking the first edge. Left: Entry and exit direction and inferred entry and exit angle memory for 7 real flies. Dashed gray line represents average of the corresponding simulations plotted for comparison. Right: Same as at left but plotted for 28 simulated trajectories.

**Extended Data Figure 10:**
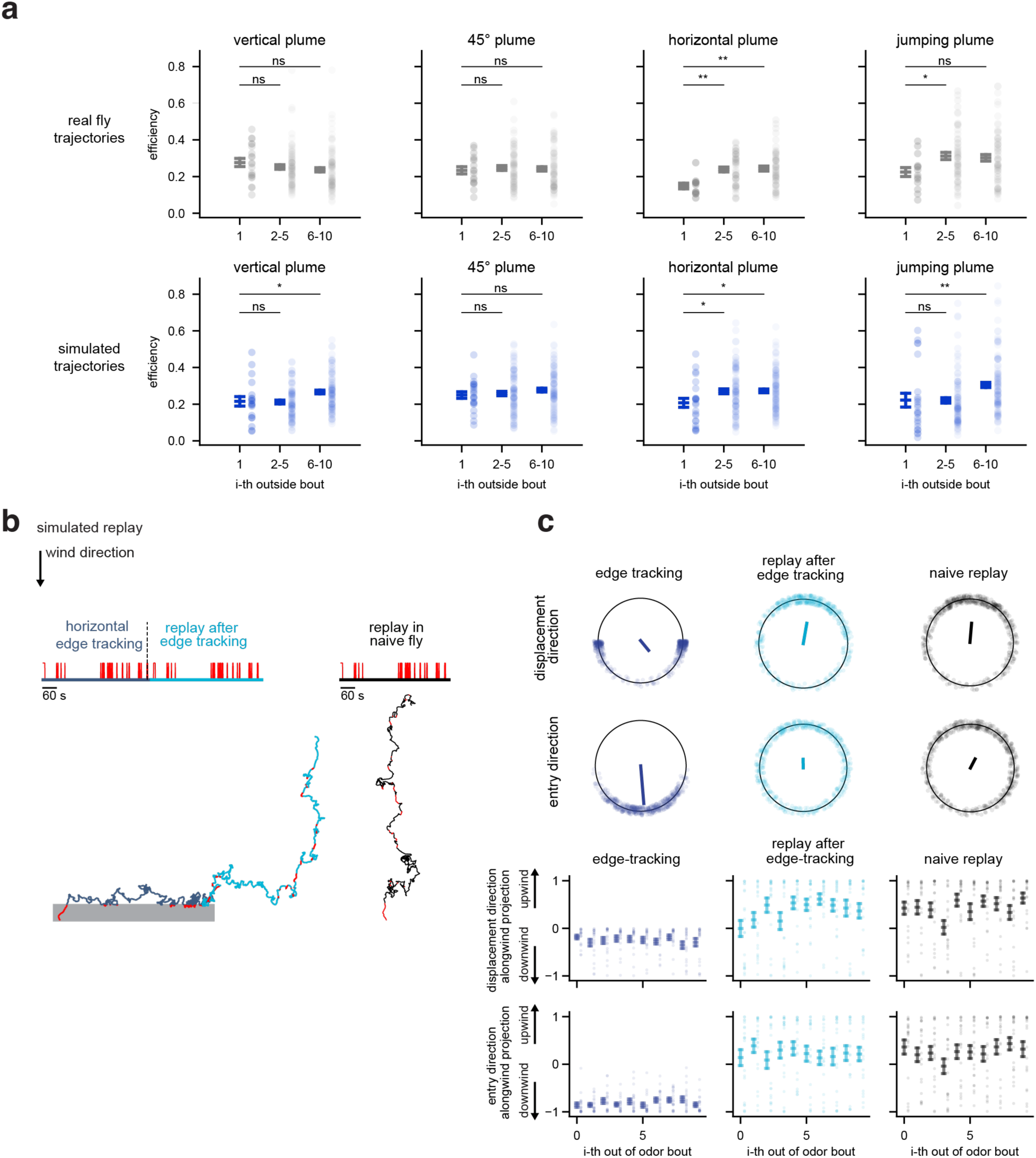
Model of how flies modify their trajectories with experience. **(a)** Plots depicting the change in efficiency of outside bouts (distance perpendicular to edge divided by total path length) for real flies (top) and simulated trajectories (bottom) tracking different plume types (0-degree, 45-degree, 90-degree, and jumping plume). n=11-20 flies; 19-23 simulations. Each point represents the efficiency of a single bout with the lines representing the average value ± standard error. **(b)** Left: Representative simulation of the olfactory ‘replay’ experiment, showing simulated trajectory during edge tracking of a 90-degree plume (dark blue trace indicates outside the odor, red inside the odor), subsequent replay of the same odor sequence (light blue trace indicates outside the odor, red inside the odor) or replay of the same odor sequence to a naïve model (black trace indicates outside the odor, red inside the odor). Right, top: Polar plots showing the net displacement direction of simulated trajectories outside the odor and entry direction when they encounter the odor during edge tracking (left, dark blue), replay (middle, cyan), or replay to naïve models (right, black). Each point represents a single bout. Solid lines represent the average vector. Right, bottom: changes of the alongwind projection of directions over outside bouts, error bars are standard error across n=21 simulations. For the alongwind projection, 1 denotes straight upwind and -1 denotes straight downwind. n.s., not significant p>0.05,* p <0.05, **p < 0.005. Details of statistical analyses and sample sizes are given in Table S1.

**Extended Data Figure 11:**
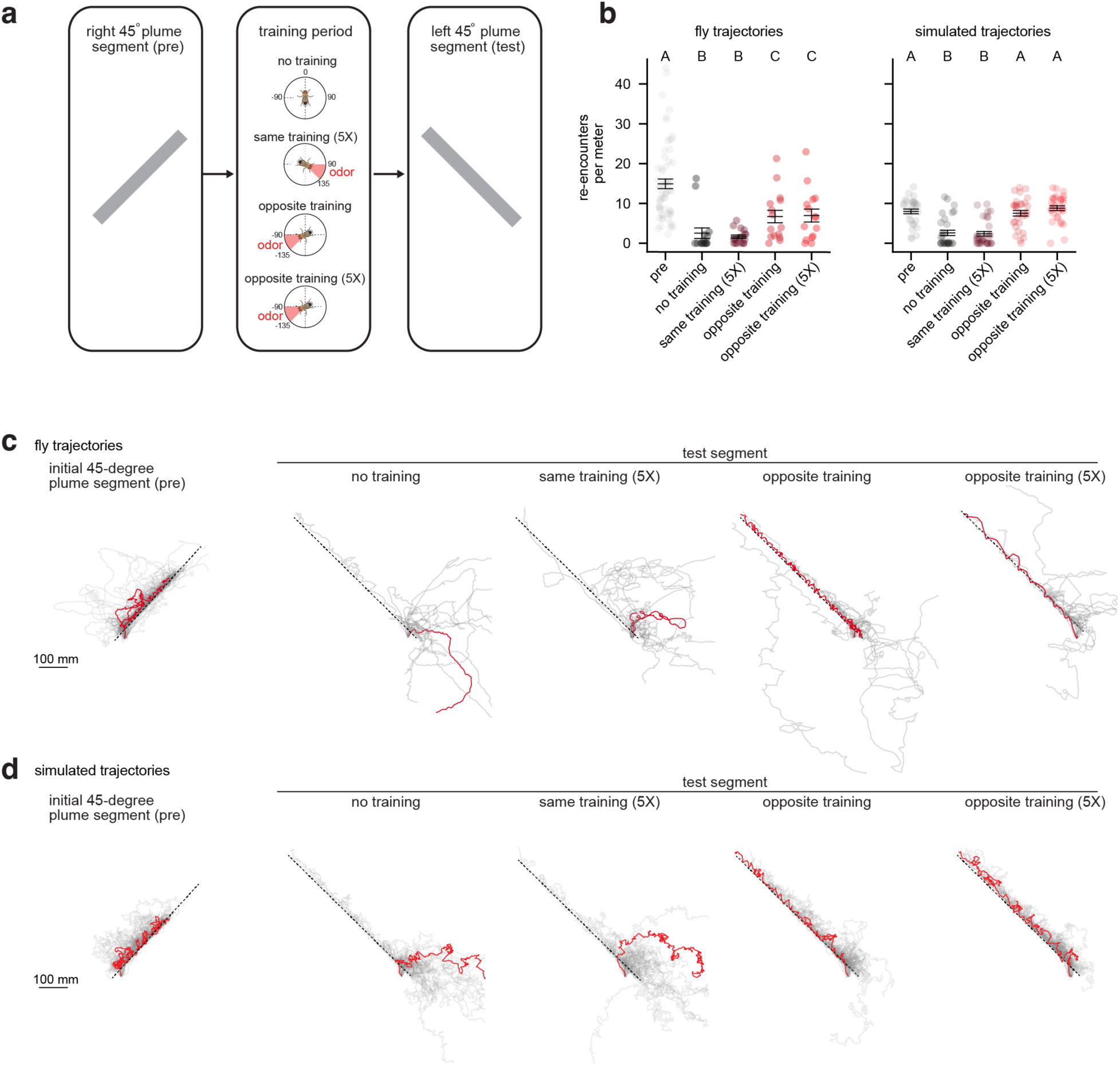
Flies can be operantly trained to track plumes of different orientations. **(a)** Schematic of experimental protocol used to operantly train flies to track plumes in different orientations. After a 5-minute baseline period during which flies walked in clean air, they were given a 45° plume to navigate. Animals that edge tracked a plume for at least 250 mm were then subjected to one of four training conditions: No training, in which flies were immediately tested with a -45° plume in the opposite direction without reinforcement; Same training (5X), in which odor delivery was triggered when flies were heading in the same direction that would lead them back to the initial plume segment (e.g. the 90° to 135° sector), and ceased upon entering a 90° upwind exit zone (-45° to 45°) (training was repeated five times); Opposite training (1X), in which odor delivery was triggered when flies traveled in the direction (e.g. -90° to -135° sector) required to enter the second plume segment. Note that to enter the second plume segment flies must travel in the same direction as used for operant conditioning (-90° to -135°) and so effectively have a second operant training epoch; Opposite training (5X), consisting of five bouts of odor reinforcement triggered by entering the -90° to -135° sector required to enter the second plume segment. Plume orientations (e.g. whether the initial plume segment was oriented 45° or -45°) were interleaved across trials for consistency and the type of trial was defined in advance. **(b)** Comparison of the plume re-encounters per meter for the five conditions for real flies (left) and simulated trajectories (right). n=17 for same training, n=15 for no training and opposite training. n=31 simulations for each condition. **(c)** Overlaid trajectories for flies tracking the indicated plume segments: initial plume, no training, same training (5X), opposite training (1X), and opposite training (5X). Individual trajectories shown as thin black line with a single representative trajectory shown in red and plume’s edge is depicted as dashed line. **(d)** Same as in (c) except for simulated trajectories. Letters denote statistically different groups (p < 0.05). Details of statistical analyses and sample sizes are given in Table S1.

**Table S1.**
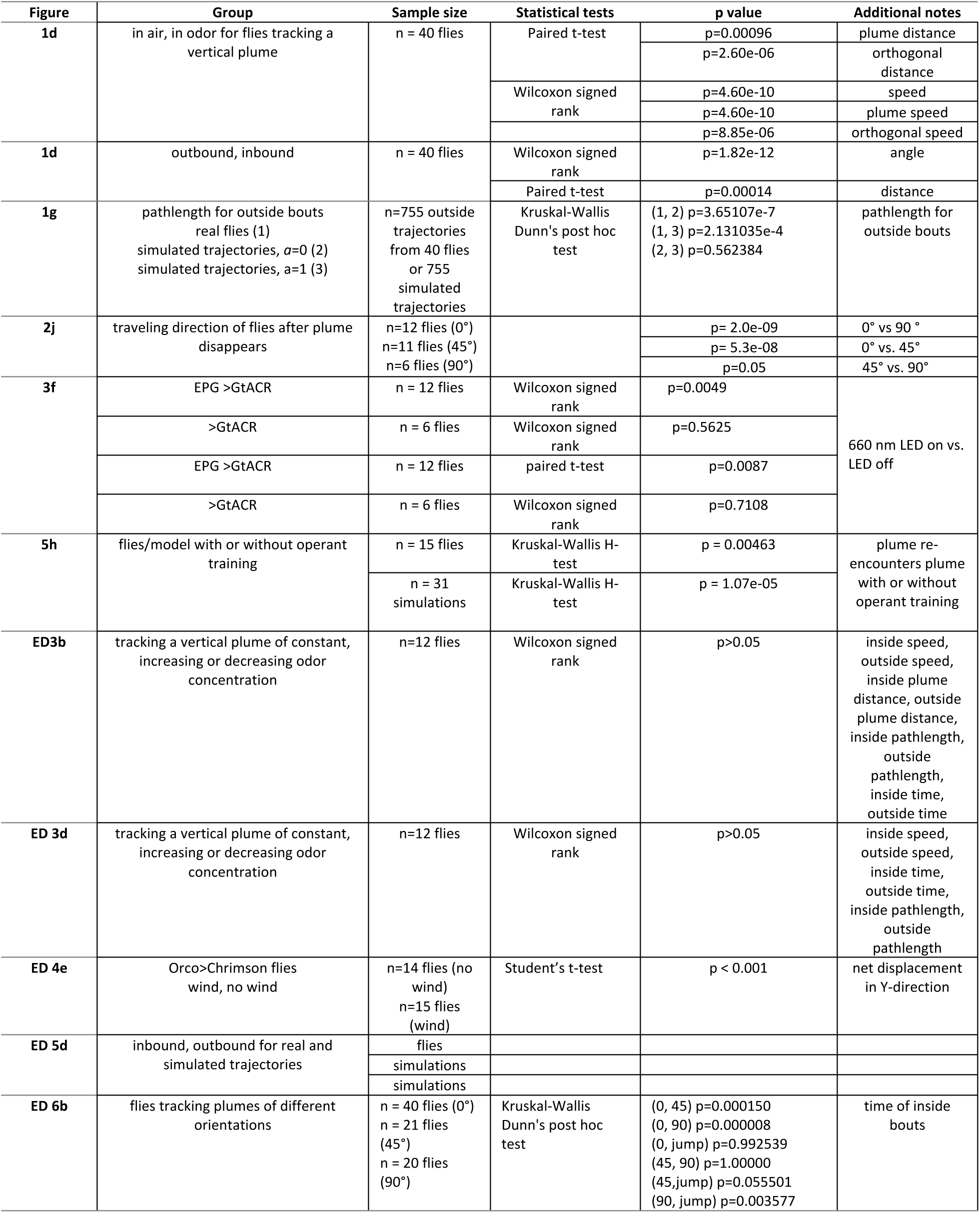
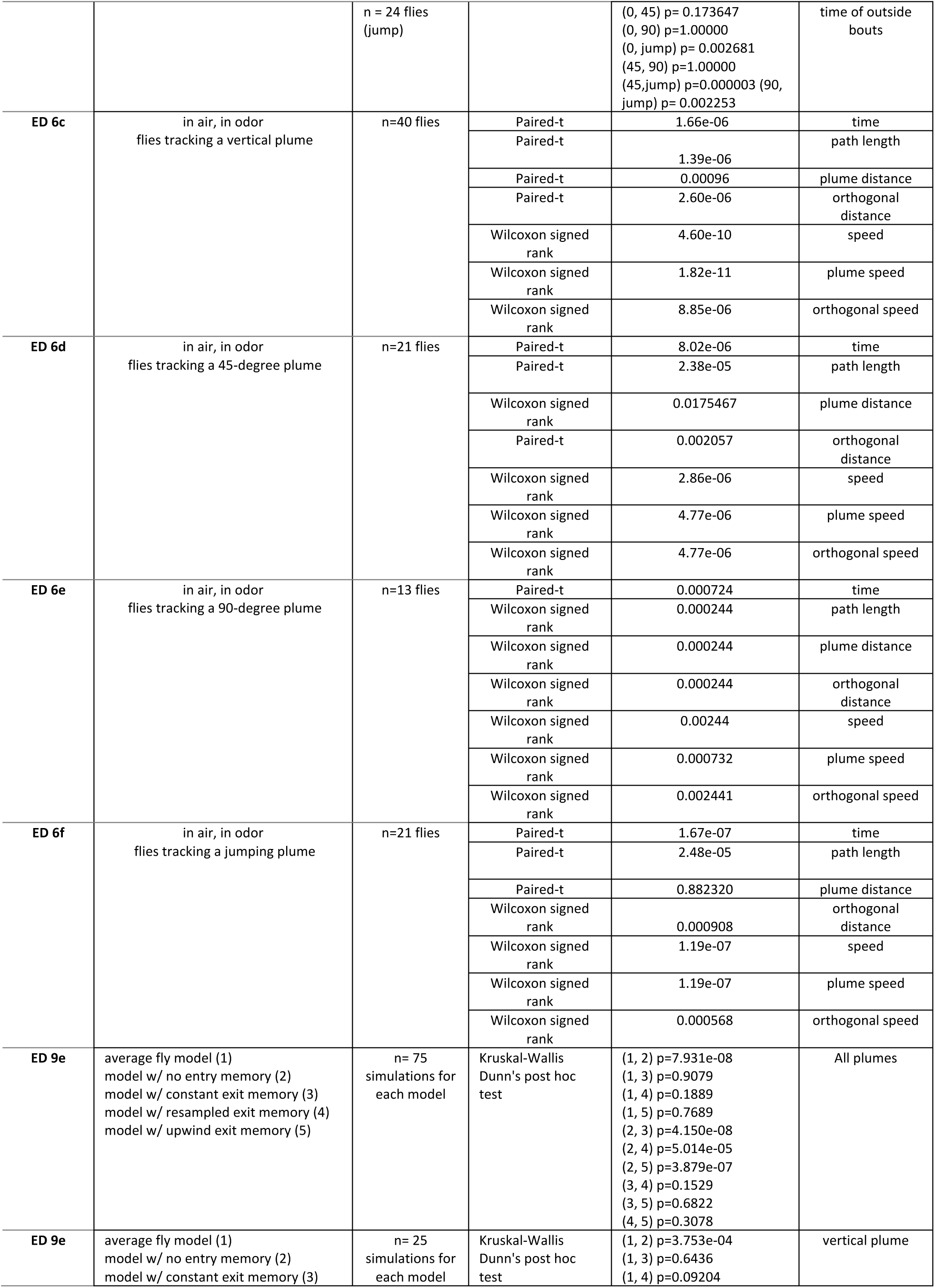
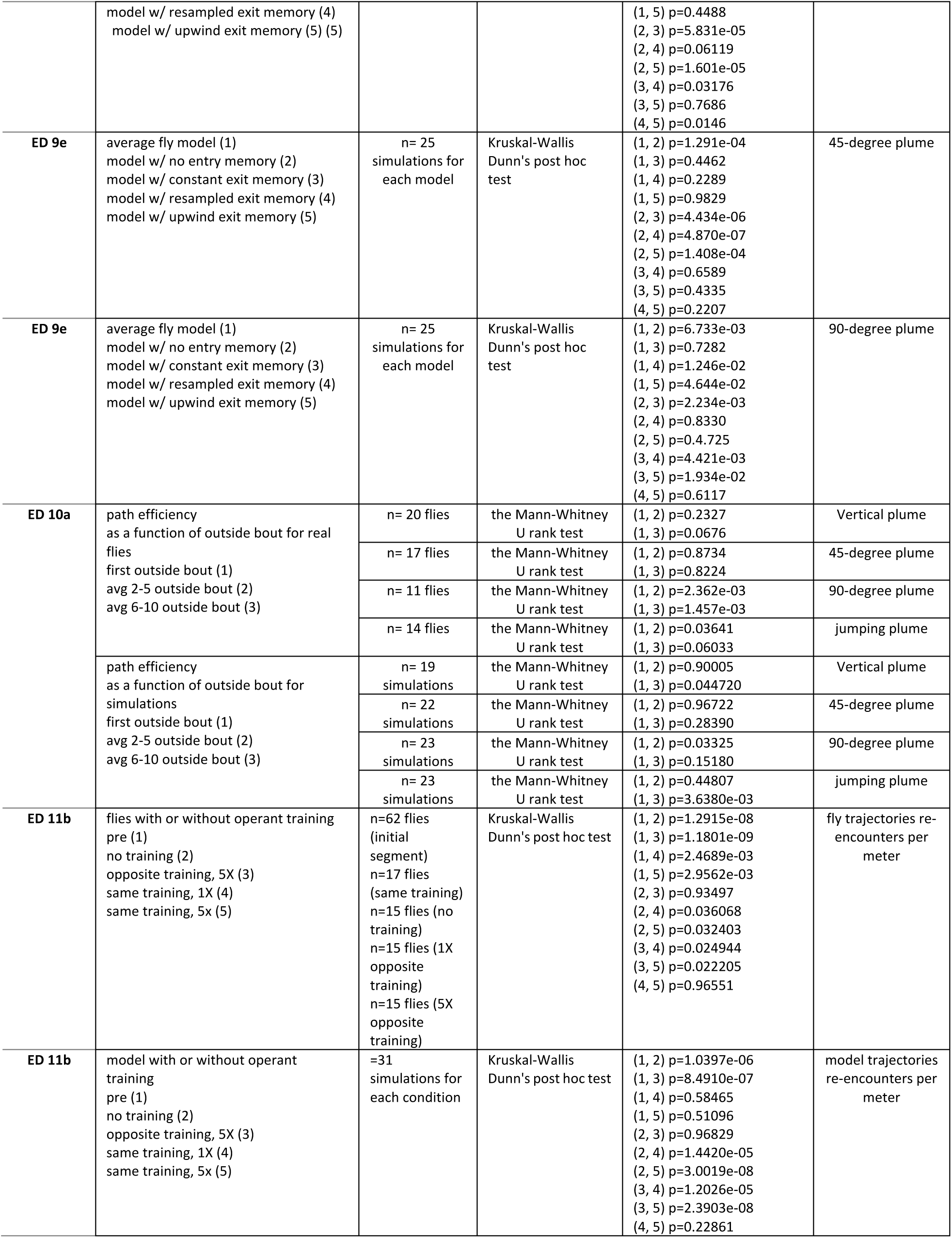
Statistics and Sample Sizes.

